# Light activation of Orange Carotenoid Protein reveals bicycle-pedal single-bond isomerization

**DOI:** 10.1101/2022.01.17.475858

**Authors:** V.U. Chukhutsina, J.M. Baxter, A. Fadini, R.M. Morgan, M.A. Pope, K. Maghlaoui, C.M. Orr, A. Wagner, J.J. van Thor

## Abstract

Orange Carotenoid protein (OCP) is the only known photoreceptor which uses carotenoid for its activation. It is found exclusively in cyanobacteria, where it functions to control light-harvesting of the photosynthetic machinery. However, the photochemical reactions and structural dynamics of this unique photosensing process are not yet resolved. We present time-resolved crystal structures at second-to-minute delays under bright illumination, capturing the early photoproduct and structures of the subsequent reaction intermediates. The first stable photoproduct shows concerted isomerization of C9’-C8’ and C7’-C6’ single bonds in the bicycle-pedal (BP) manner and structural changes in the N-terminal domain with minute timescale kinetics. These are followed by a thermally-driven recovery of the BP isomer to the dark state carotenoid configuration. Structural changes propagate to the C-terminal domain, resulting, at later time, in the H-bond rupture of the carotenoid keto group with protein residues. The isomerization and its transient nature are confirmed in OCP crystals and solution by FTIR and UV/Vis spectroscopy. This study reveals the single bond isomerization of the carotenoid in the BP manner and subsequent thermal structural reactions as the basis of OCP photoreception. Understanding and potentially controlling the OCP dynamics offers the prospect of novel applications in biomass engineering as well as in optogenetics and bioimaging.

## Main text

Carotenoids are found in plants, algae, and microorganisms and constitute the most widely spread group of pigments^1^. Carotenoids are isoprenoids made-up of a central carbon chain of alternating single and double bonds with unique photochemical properties. The S0-S1 electronic transition is optically forbidden in carotenoids and light absorption occurs through the intense S0-S2 transition^2^ which is responsible for their yellow, orange and red colours. After light absorption, the S1 dark state is populated through internal conversion, which then relaxes with a picosecond lifetime^3^. The structure and conformation of carotenoids determine the energy landscape of their electronic states and, as a result, their biological functions. Carotenoid chromophores function as key photosynthetic regulators of biomass production^4,5^ when their S1 states act as efficient quenchers of chlorophyll excited states^6,7^. Furthermore, carotenoid excited states also act as energy donors in photosynthetic light-harvesting^3^ as well as photoreceptors in light-signalling^8^. The structural and molecular basis for the diverse biological functions of carotenoid chromophores is of high importance and is necessary to fully understand in order to gain control of these processes. The carotenoid canthaxanthin (CAN) bound to the Orange Carotenoid Protein^9,10^ is responsible for a light signalling function. OCP undergoes a photocycle^9^ in which the keto-carotenoid also plays several different roles: in the inactive or orange state (OCP^O^, Fig 1a), it senses blue-green light to initiate its photocycle, resulting in a final product that has undergone substantial structural conformational changes and exhibits distinct separation of the two OCP domains, namely, the all-helical N-terminal domain (NTD) and an α/β-fold C-terminal domain (CTD)^11^. In the active state, following OCP^R^ (red state) binding to the cyanobacterial light-harvesting antenna, the keto-carotenoid dissipates excess absorbed light energy through non-photochemical quenching (NPQ), a vital process for cyanobacterial survival in turbulent waters^12,15^. In cyanobacteria NPQ fully builds-up over minutes upon OCP control^12,13^. Therefore, resolving OCP photocycle is essential for understanding and controlling OCP photoregulation.

**Fig. 1:**
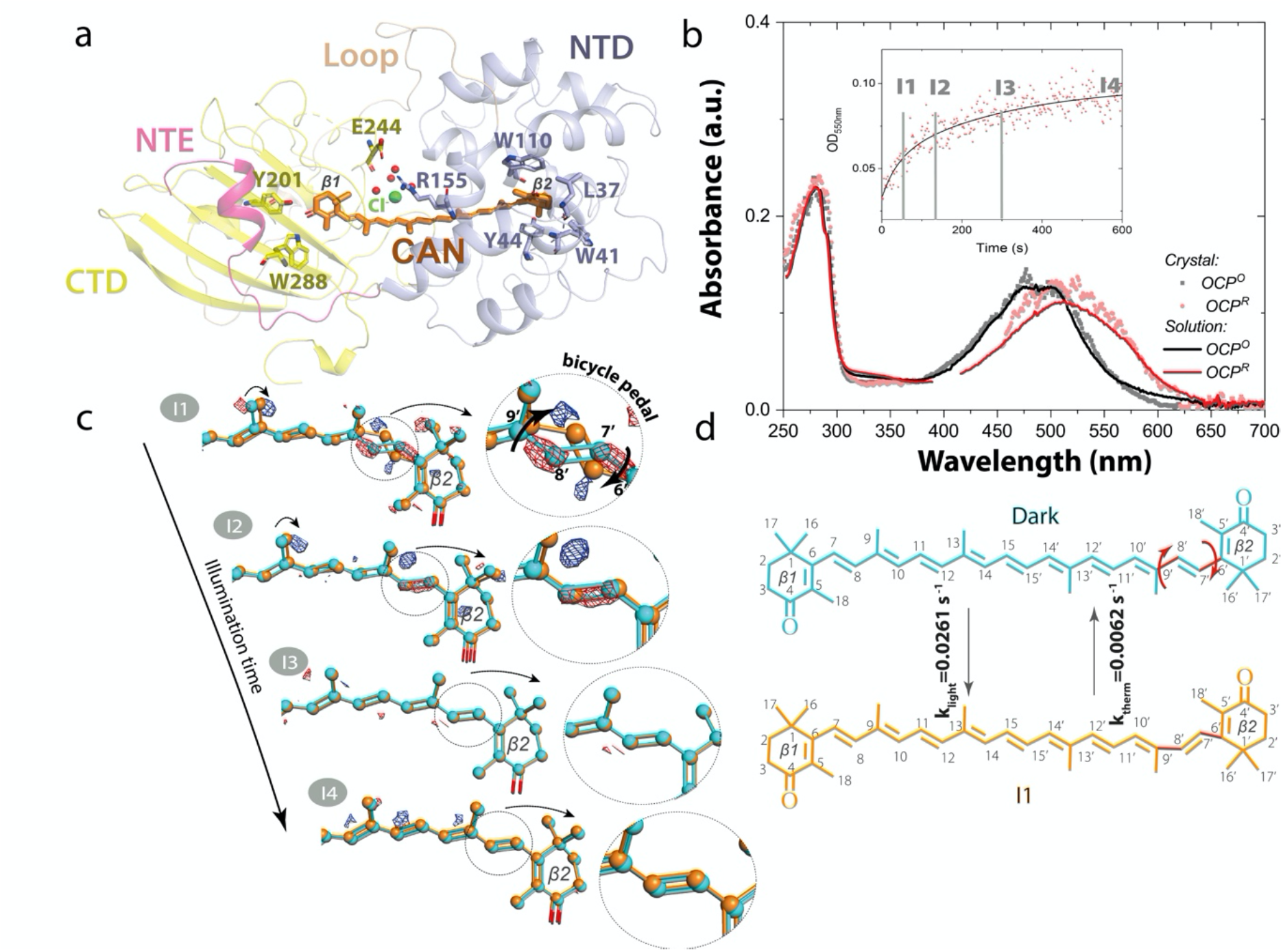
Canthaxanthin single bond isomerization in the bicycle-pedal manner triggers OCP light activation. (a) Structure of OCP in its inactive state (OCP^O^) displaying the C-terminal domain (CTD, yellow colour), N-terminal domain (NTD, violent colour), N-terminal extension (NTE, pink colour) and loop (wheat colour). The canthaxanthin (CAN) carotenoid is embedded between the two domains. Cl^—^ (green sphere) is resolved in the CTD where it coordinates four water molecules (red spheres) through ion-dipole interactions at distances of 3.0-4.0 Å. The closest CAN carbon (C12) is at 4.1 Å distance (Extended Data Fig 1). (b) Absorbance changes of OCP in solution (solid lines) and crystalline state (dotted lines) upon illumination with violet light (at 410 nm with 3.5 mWcm^-2^ power; FWHM 20 nm of the emission spectrum). Inactive state OCP^O^ and active state OCP^R^ are shown in black/gray and red/pink respectively. The OD intensities are unavoidably polluted by the LED emission in 380-430 nm and therefore were omitted from the graph. Insert: Absorbance changes (pink dots) in the crystalline state detected at 550 nm upon 10 min of illumination. Black line represents double exponential fit of the curve (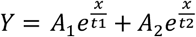, where *A*_*1*_ =0.02, *t*_*1*_ =38 s and *A*_*2*_ =0.44, *t*_*2*_ = 291 s. Grey labels indicate time points when I1-I4 intermediates have been cryotrapped(c) Difference electron density (DED) maps were obtained for the four states: (I1) 1 min (F_I1_ – F_Dark_), (I2) 2 min (F_I2_ – F_Dark_), (I3) 5 min (F_I3_ – F_Dark_), (I4)10 min (F_I4_ – F_Dark_). Red: negative DED, blue: positive DED on the −3σ/3σ contour levels, respectively. Carotenoid coordinates of dark (OCP^O^) and light states (I1-I4) are represented by cyan and orange lines respectively. Illuminated coordinates were obtained from extrapolated electron densities. Populations of the illuminated states obtained from the occupancy refinement (see Materials and Methods) are 15% (I1), 22 % (I2), 25 % (I3) and 32% (I4). Arrow in the insert in l1 state indicates carotenoid movement during illumination. (d) CAN in C9’-C8’ *trans*/ C7’-C6’ *cis state* is present in OCP^O^ while C9’-C8’ *cis*/ C7’-C6’ *trans* conformation (bicycle pedal isomerization) is observed in I1. The latter thermally recovers back to OCP^O^ CAN state with prolonged illumination time (after 2-5 min).

The mechanism and primary photochemical reaction of the keto-carotenoid that triggers this OCP^O^-OCP^R^ conversion is currently debated and, therefore, requires structural evidence^8,14^. It has been argued that OCP photoactivation is initiated by H-bond rupture between the 4-keto group of the β1-ring and neighbouring residues Y201 and W288.^15–21^ Several different proposals have been made for the primary photochemistry resulting in this particular H-bond rupture: (1) a charge transfer state^22^; (2) a shift in keto–enol equilibrium^18^; (3) a transfer of vibrational energy from the carotenoid to the protein leading to the H-bond rupture^23^, similar to H-bond rupture in bacteriorhodopsin ^24^; (4) a structurally distorted form of the carotenoid in the S1 state characterized by increased carotenoid planarity^19^; (5) a photoinduced s-isomerization of the C6−C7 single bond, flipping the orientation of the β1-ring by 90°^15^. Support for s-isomerization in OCP stemmed from the X-ray structure of the isolated NTD (also known as red carotenoid protein, RCP), in which the keto-carotenoid was present in a C6−C7 s-*cis* conformation^15^, and frequency calculations showing agreement between a C6−C7 s-*cis* conformation and OCP^R^ Resonance Raman data^22^. However, the hypothesis for s-isomerization in OCP^R^ is not consistent with recent infra-red (IR) anisotropy observations^19^. A recent study of the fluorescence spectra and yields of OCP^O^ with different carotenoids suggested that the mechanism of OCP photoactivation does not involve the keto group of the β1 ring but includes out-of-plane motions of the β2 ring^25^. On the other hand, in the closest to OCP photoreceptor, rhodopsin, its photosensing pigment, retina, which is actually synthesized from a carotenoid by irreversible oxidative cleavage, undergoes bicycle pedal (BP) double pond rotation as part of vision photokinetics. ^26^ However, never such a photoresponse has been proposed in OCP. Here we show time resolved crystallographic and spectroscopic evidence which resolve the initial photoproducts and subsequent structural propagation in the activation process.

Figure 1b shows the light induced absorption changes measured in crystals and solutions of OCP. In contrast to a previous crystallographic investigation of OCP photoactivation^18^, we observe that the absorbance spectra of OCP^O^ in solution and in crystals are nearly identical (Fig 1b), thereby providing evidence for the same electronic structure of the carotenoid in both crystals and solution. Illumination of crystals with 410 nm light and 3.5 mWcm^-2^ power density causes a significant spectral red-shift of 25 nm due to OCP^O^-OCP^R^ conversion within 5 min on average (Fig 1b insert). In solution similar spectral shift is observed. It occurs on the same timescale as in crystals, but slightly faster (3 min) as resolved from the detailed spectroscopic analysis described later in the manuscript. All in all, strong similarity of photoresponses in the crystals and in solution allow us to conclude that OCP is photoactive in crystals with the same photoresponse characteristic as seen in solution. An early reaction intermediate was captured by intense illumination and cryo-trapping of OCP crystals. The crystal structure and carotenoid conformation was determined at 1.3 Å resolution for the dark state (OCP^O^, Fig 1a) and was found to be in good agreement with those reported in previous X-ray studies^9,15,16,18^ (Extended Data Table 1).

The data furthermore supports identification of glycerol, acetate, Cl^-^ and 401 water molecules in the crystal structure (Extended Data Fig 1, Extended Data Table 2-3). The location of Cl^-^ in this study was confirmed by long-wavelength X-ray crystallography (Extended Data Fig 1, Extended Data Table 3). In brief, the contribution of Cl^-^ to the anomalous scattering signal at 2.9keV is 3.95 e^-^ (f’’), whereas at 2.8keV, this value is 0.40 e^-^ (f’’). The difference between these two values allows for specific sites of Cl to be identified by observing the anomalous difference Fourier map at the expected Cl site for each (Extended Data Fig 1b).

Cl^-^ is resolved in the CTD, where it coordinates four water molecules through ion-dipole interactions at distances of 3.0-3.2 Å (HOH16, HOH23, HOH46) and 4.0 Å (HOH41). The closest CAN carbon is C12 located at 4.1 Å distance from Cl^-^ in CTD (Extended Data Fig 1a). Three amino acids (L248, T275, P276) were resolved within 4 Å from Cl ^-^. The electron density of the resolved Cl^-^, was assigned to Cl ^-^ in the first OCP crystal structure (PDB entry 5UI2) based on the signal intensity and crystallization conditions ^9^, but subsequently has been assigned to a water molecule in several reports: HOH609 (PDB entry 5TV0), HOH699 (PDB entry 5TUX), HOH673 (PDB entry 4XB5) and HOH1020 (PDB entry 3MG1).

In addition to OCP^O^, X-ray crystallography data was collected of four illuminated structural intermediates I1, I2, I3 and I4 corresponding to 1 min, 2 min, 5 min and 10 min of illumination. Data collection and refinement statistics were comparable to that of the OCP^O^ reference data. Q-weighted F_Light_–F_Dark_ difference electron density (DED) maps^27^, indicating light-induced changes in X-ray data, and CAN coordinates at different time points(see Materials and Methods, Extended Data Fig 2) are presented in Fig 1c.These illuminated coordinates (I1-I4, Fig 1c) were resolved through their refinement to extrapolated structure factors based on the commonly used approach described before^28^. The I1 dataset with 1 minute illumination reveals s-trans to s-cis CAN isomerization along the C9’-C8’ single bond concerted with s-cis to s-trans isomerization along the C7’-C6’ single bond (BP isomerization) as the early step of activation (Fig 1 c-d, Extended Data Fig 3 a). As a result, C8’=C7’ changes its orientation (Fig 1c-d). The isomerization-related DED signals are also detected in I2, though less pronounced compared to I1. In contrast to I1 the resolved extrapolated coordinates of I2 support only a minor population of the concerted C9’-C8’ *cis*/ C7’-C6’ *trans* state while the dark (C9’-C8’ *trans*/ C7’-C6’ *cis*) conformation becomes again the dominant state. The crystallographic observations for the first 2 min of illumination go in line with the kinetics and spectra determined by solutions spectroscopy described below and identify the bicycle pedal (BP) ^29^ isomerization. Unlike the BP mechanism, driving the first steps of vision in rhodopsin retina, the BP photoresponse observed here is novel involving single bond isomerization and has never been reported before. It occurs with k_isom_=0.0263 s^-1^ and is followed by thermal relaxation (k_therm_=0.0062 s^-1^, Fig 1 d). The isomerization fully decays after 5 min in I3 and I4 judging from DED maps, 2mFo-DFc maps, and refined coordinates using extrapolated structure factors (Fig 1c, Extended Data Table 1).

While resolved extrapolated coordinates of CAN in I1-I4 states identify the CAN BP isomerization within 1 min of illumination at the experimental light intensity used at ambient temperature as a first light-activation step, UV/Vis spectroscopy confirms that the rate of its accumulation is proportional to the illumination intensity (Extended Data Fig 4 a-c). Complete thermal relaxation of CAN back to dark conformation occurs after 5 min illumination but also in darkness (Extended Data Fig 4 a, Supplementary information) indicating that the process is thermally-driven. It goes in line with the fact that single bond rotation in conjugated systems does not require pi-bond breakage, therefore could occur at ambient temperatures. In contrast to the current prevailing model ^15–21^, which proposes a H-bond rupture in CTD as the primary photoproduct of the OCP photocycle, our X-ray data directly demonstrate that a transient BP ^29^ single bond isomerization is generated prior to rearrangements in CTD. This finding reveals the molecular activation mechanism that precedes the formation of product states described in the literature. The I1 structure is a first example where BP ^29^ isomerization of a carotenoid carbon chain regulates the activity or the biological function of a protein. We cannot exclude that the isomerization of the two single bonds as seen in I1 occurs sequentially within 1 min, however it is very unlikely, since BP process is a space-saving process, therefore was demonstrated to be favorable in a tight-protein cavity ^26,30^ or other restraining environment^31^.

Photoinduced structural changes occur throughout the OCP protein and propagate in a time-dependent manner. The F_I1_–F_Dark_ map reproducibly displays differences in three regions (Fig 2, Extended Data Fig 2): in the NTD with the highest DED amplitudes within 6 Å from the CAN β2 ring (Zone 1, Fig 2) and in the N-terminal extension (Zone 2, Fig 2). The DED signals appear in the CTD (Zone 3, Fig 2, Extended Data Fig 2) with time delays longer than 1 min. In the I2 intermediate (i.e. after 2 min of illumination), the DED signals appear primarily in the region of the CAN binding pocket and subsequently appear along helices of the CTD in the I3 and I4 structures (Fig 2). Concomitant with the development of DED signals in the CTD, the DED signals in Zones 1 and 2 decay in I3 and I4. These results indicate that NTD is the primary site of action in the OCP light-activation. Therefore, the initiation of photo-sensing in the OCP photocycle involves structural changes in the NTD rather than in the CTD as previously suggested^18^. B-factor analysis on the whole protein level (Fig 2) and on the level of CAN (Extended Data Fig 3c) show that B factors is accompanied with appearance of substantial structural changes in these states and/or areas.

**Fig. 2:**
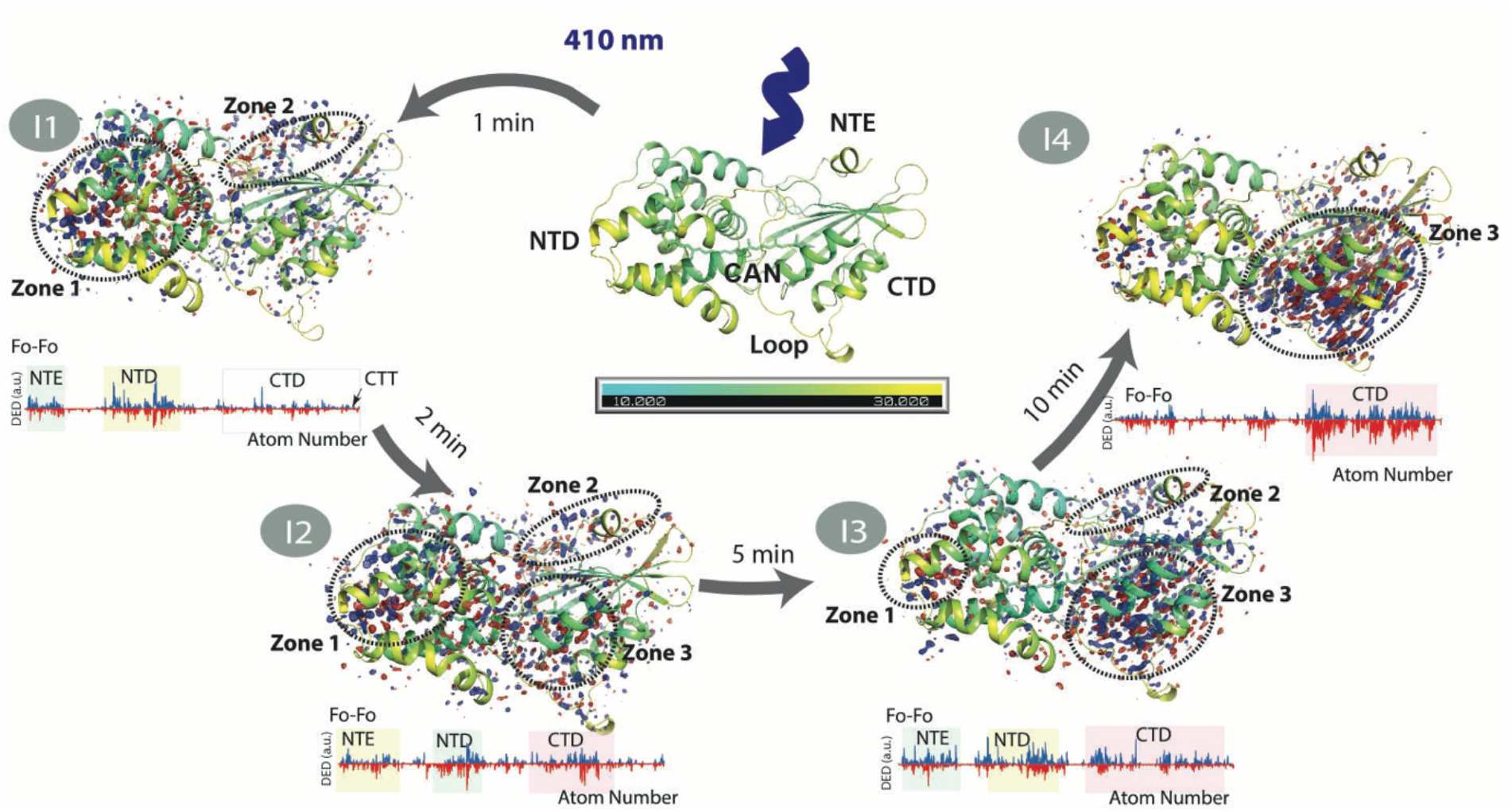
Overall structural effect of illumination on OCP structure. Datasets from which all images are derived (Dark, I1-I4) have a resolution of 1.3 -1.4 Å (Supplementary Information) and time points at which crystals were cryo-trapped are indicated for I1-I4. Each structure is coloured in cyan-yellow gradient based on B-factor (10.0(cyan)-30.0 (yellow)). (I1-I4, upper panel) F_0_-F_0_ Difference electron density (DED) maps (blue/red at ±3.5σ) for the four states shown below each corresponding structure: (I1) 1 min (F_I1_– F_Dark_), (I2) 2 min (F_I2_– F_Dark_), (I3) 5 min (F_I3_– F_Dark_) and (I4) 10 min (F_I4_– F_Dark_). The DED signals are localized in the 3 major areas: *zone 1*: NTD; *zone 2*: NTE; *zone 3*: Alpha-helices in CTD. Note that some DED signals are observed at the interface between the two domains (red arrows in I1 and I2) while the C-terminal tail (CTT) does not exhibit any noticeable DED signal. Red: negative DED, blue: positive DED on the −3.5σ/3.5σ contour levels, respectively. See main text for explanations. (I1-I4, lower panel) DED signals are also shown as the sequential integrated signals^47^ around each atom in the protein chain. The figure was created using PyMOL Molecular Graphics System (Version 2.5.0).

The structural rearrangements in the intermediates I1, I2, I3 and I4 are analysed in detail following coordinate refinement of extrapolated structure factors. Tyrosine Y44 and tryptophan W110 are located in closest proximity (4 Å) to the C6’-C7’=C8’-C9’ moiety in the hydrophobic pocket which also contains the β2-ring of the carotenoid (Fig 3 a). Together with W41, residues Y44 and W110 interact with the carotenoid β2-ring through *π*-*π* stacking (Fig 1 a).^16^ In the I1 structure, no changes of coordinates are observed for W110 compared to OCP^O^ but the Y44 sidechain undergoes a rotation by 31° (Fig 1a). This observation suggests that the Y44 sidechain reacts to the BP isomerization to trigger structural changes in NTD. The structures of the I1 and I2 intermediates suggest that the repositioning of the Y44 sidechain causes the displacement of multiple amino acids in NTD. These, among many others, include residues Y111 and G114, which are closest to Y44 and are displaced in I1 and I2 (Fig 3a). In agreement with our crystallographic observations, previous spectroscopic analysis demonstrated that the Y44S mutation fully inhibits OCP^O^-OCP^R^ conversion^21^. As a result of Y44 rotation by 31° in NTD in I1 state, the hydroxyl group of Y44 gets 2.0 Å closer to C9’ carbon, substantially redistributing charge-density around C9’-C8’ (Extended Data Fig 4 d-e). These changes are transient and the amino acids return to their original location as in OCP^O^ after 5-10 min of continuous illumination (Fig 3 a). Leucine L37, which has been proposed to play an important role in OCP light-activation ^25^, does not undergo any significant movement upon illumination (Fig 3a). We therefore conclude that the BP isomerization of CAN in I1 causes rearrangement of the CAN β2-ring binding pocket in the NTD, which is mediated by contact with Y44.

**Fig. 3:**
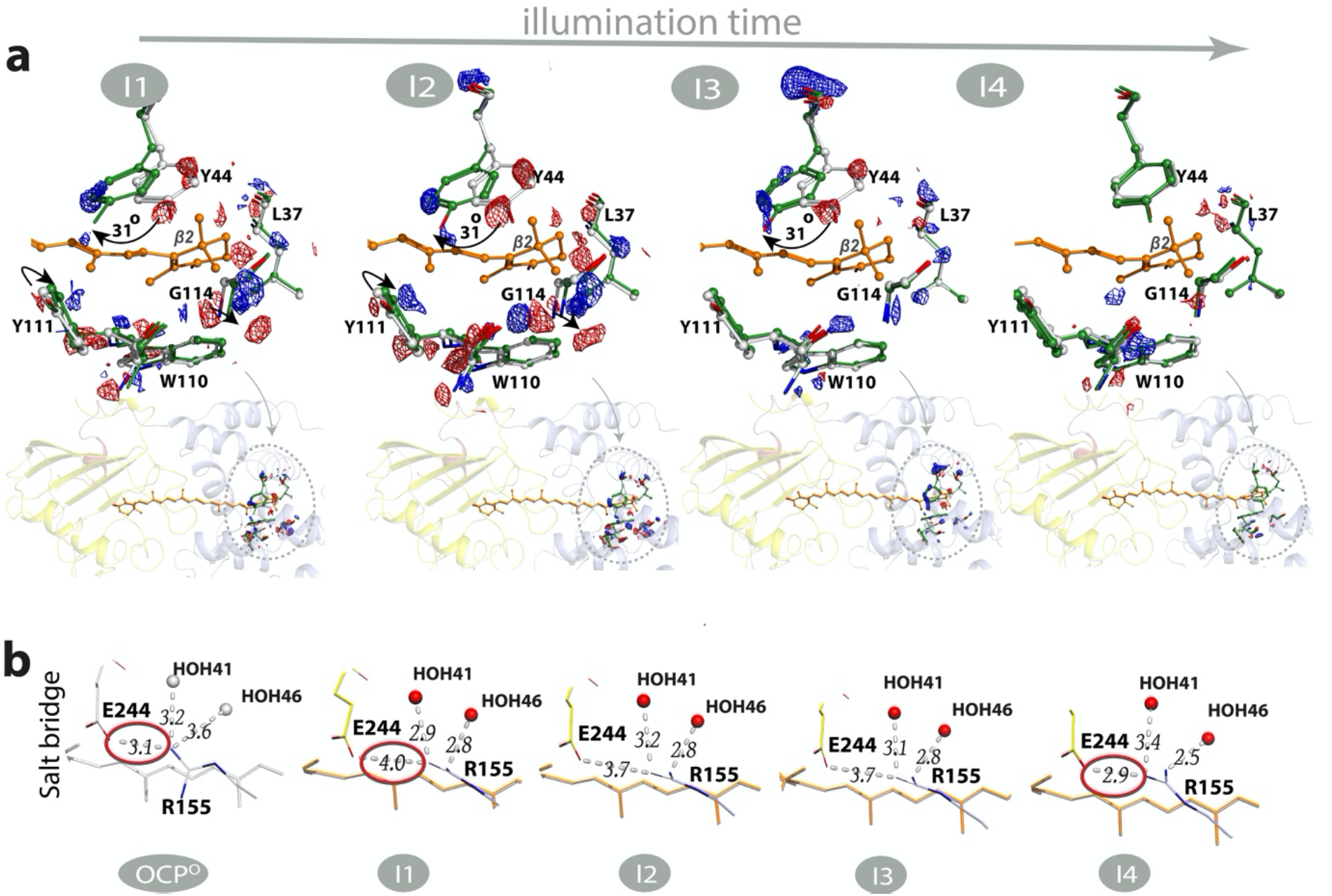
OCP structural changes in I1-I4 states upon illumination. (a) Dark (OCP^O^) coordinates are represented by grey sticks while light state coordinates obtained from extrapolated maps (I1-I4) are represented by green (NTD) and orange (CAN) sticks. Difference electron density (DED) F_Light_–F_Dark_ are shown with blue/red mesh at ±3.0σ). (b) Changes in the salt bridge formed between residues E244 and R155. The figure was created using PyMOL Molecular Graphics System (Version 2.5.0).

The rearrangement of the β2-ring binding pocket coincides with further conformational changes within the NTE and the NTD. This affects primarily helices αA-α C and αE - αH, as can be judged from the location of DED signals (Fig 2). The amplitude of these signals is maximal after a 1 min illumination (F_I1_–F_Dark_) followed by a decrease after a 2 min illumination (F_I3_–F_Dark_) and complete decay after a 10 min illumination (F_I4_–F_Dark_). The conformational rearrangements of the αA-αC and the αE - αH helices affect the hydrogen-bond network by increasing the number of polar contacts in the main backbone of I1 by 5% compared to OCP^O^. In I1, 14 new H-bonds are created in the backbone which gradually decay in I2-I4 (Extended Data Table 4). These results indicate that, following light-activation, the light-sensing process in OCP is mediated not only by the CAN β2-ring binding pocket but involves the hydrogen-bonding of the whole domain.

The DED signals within a 5 Å radius from the carotenoid at the CTD and NTD interface (Fig 2) are assigned to the modification of the salt bridge between R155, E244 (Fig 3b) and the reorganisation of water molecules (Fig 3b, Extended Data Fig 5). The salt bridge rupture between R155 and E244 is already observed in I1 and it is noticeable as a increase of the distance between residues E244 and R155 by 1 Å (Fig 3b). The position of R155 is affected by the water molecules HOH41 and HOH46 through H-bonding (Fig 3b, Extended Data Fig 6 a). The positions of Cl^-^ anion, HOH41, HOH46, and other water molecules in the water channels are rearranged during the illumination period (Extended Data Fig 5-6, Fig 3b). Overall, the movement of Cl^-^ anion observed in the first 5 min of illumination (I1-I3) is mainly caused by the H-bond rearrangement in the water channel (Cl1 in Extended Data Fig 5, Extended Data Fig 6 a and Supplementary information), while in I4 it is mainly caused by translocation of the CTD away from NTD (Extended Data Fig 6 b).

In the final illumination product (I4, Fig 3b) the distance between residues E244 and R155 is restored to the original 3 Å distance found in the dark state, suggesting that, although an important event in the initial steps of light activation, salt-bridge rupture may not be a prerequisite for subsequent domain separation. Previously reported FTIR and X-ray protein footprinting results suggested that signal propagation of OCP conformational change is mediated by rearrangement of internal hydrogen bonding networks^11,32^. We complement this knowledge by showing that salt bridge rupture between R155 and E244, the important precursor of NTD and CTD separation, occurs in the first illumination step (I1) and is controlled by a hydrogen bonding network involving protein-bound water molecules.

In addition, an increase in DED signal intensities in the CTD is observed with increasing illumination time (Zone 3, Fig 2). This is first manifested in CTD amino acids L248 and T275 among others located at the interface of CTD and NTD (Extended Data Fig 6 a). The DED signals for the CTD (Fig 2) do not represent a complete domain rearrangement but rather a rigid-body motion of the CTD domain in the direction away from the NTD and the dimer interface (Extended Data Fig 6 b) similar to that previously reported^18^. This CTD translocation, which was previously observed and seems to be a prerequisite for H-bond disruption^18^, is only observed at longer illumination times (after 5-10 min, Extended Data Fig 6) and only after rearrangement of the NTD binding pocket. Therefore, the H-bond rupture represents a secondary effect caused by domain separation and it is not a photo-driven process as previously proposed ^18,19^.

Next, we performed FTIR and UV/Vis spectroscopy in solution using the same illumination conditions as for crystals in order to confirm that CAN isomerization is photoinduced (Fig 4) and determine whether the same photoactivation reactions observed in crystals also take place in solution. Fig 4 displays the results from the combined global analysis fits of the UV/Vis and FTIR data using a well-established approach to analyse time-resolved spectroscopic data, which yields Evolution-Associated Spectra (EAS) components^33–35^. Since the experiment in this study was performed under continuous illumination, the EAS components do not represent intrinsic lifetimes of the intermediate states but rather dissected population accumulation rates at given illumination power densities (Extended Data Fig 4 a-c). Four components of spectral regions (EAS1-4) are identified to describe the absorbance changes in both UV/Vis and IR spectra with lifetimes of 13 s, 38 s, 162 s and a steady-state component (infinity). Note that the time acquisition rate of FTIR is below the EAS1 risetime, EAS1 FTIR spectra cannot be resolved. The concentration profiles of EAS components in time are presented in Fig 4c, insert.

**Fig. 4:**
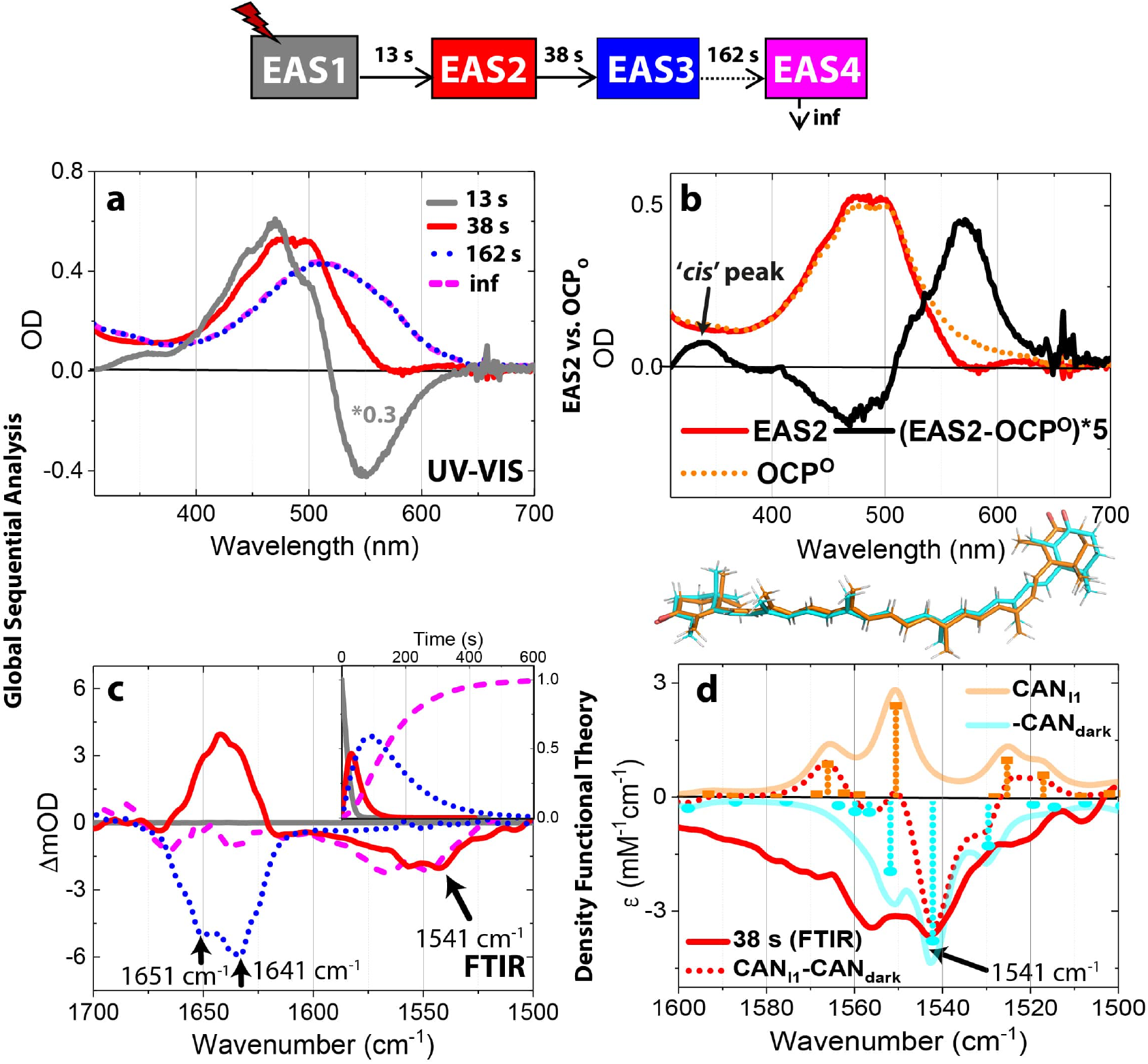
Spectroscopic evidence for bicycle pedal isomerization in OCP in solution. (a, c). Results of the simultaneous global fit of UV-VIS (a) and FTIR (c) data upon 10 min of illumination. The kinetic model used for the fit is shown on top of the panel with the respective lifetime for each transition between intermediates. The same colour code is used for the corresponding EAS components in panels a and c. Refer to the main text for an explanation of the relationship between the resolved EAS and the illuminated structural intermediates I1-I4 (Fig 1c and Fig 3). (a) UV-VIS EAS. For EAS1 spectrum multiplication factor 0.3 was used.(b) EAS2 component was compared to the original OCP^O^ spectrum. (c) FTIR EAS (in D_2_O); FTIR (in H_2_O) is shown in Extended Data Fig 4 f. Insert: Relative concentration profiles for each EAS. (d) Theoretical IR frequencies and intensities were calculated using Harmonic frequency calculations (HFC) at the B3LYP/6-311+G level. IR values were calculated for CAN in the two (CAN_Dark_ and CAN_I1_) states using CAN coordinates extracted from the extrapolated maps. Theoretical IR frequencies and calculated FTIR spectra are presented by sticks and solid lines respectively. Optimized CAN coordinates from HFC are shown on top of the panel. CAN_I1_-CAN_Dark_ (red dotted line) obtained from HFC agree with the 1542 cm^-1^ bleach from the 38-s EAS2 (red solid line).

The overall photocycle kinetics in OCP solution and OCP crystals seems to be very similar (Fig 1b, Fig 4 a and justification below). Therefore, the contribution of each component (EAS) in I1-I4 states should be approximately as follows: (i) I1 state: EAS1 (3%), EAS2 (40%) and EAS3 (57%), (ii) I2 –EAS2 (13%) and EAS3 (87%), (iii) I3 –EAS2 (2%) and EAS3 (98%) and (iv) I4-EAS3 -100%. Note that the absence of the final component EAS4 in crystals is probably due to crystal packing constraints on side-chain conformational dynamics so it was excluded from the population estimates (see the justification below). The first component (EAS1) exhibits a positive and a negative spectral signature in the visible region (Fig 4a, grey solid line) peaking at 475 nm and 550 nm respectively. We assign this spectrum to the equilibration process between OCP^O^ and the newrly forming intermediate species EAS2. The newly formed species presented by EAS2 (at 38s) has a very similar visible spectrum to OCP^O^ (t_0_ spectra) though like OCP^R^ it has more red shifted absorption peaking at 575 nm, which goes at expense of the 470-nm absorption decrease (Fig 4b, black line). But unlike in OCP^R^, the red shift of the spectrum is much less pronounced in EAS2. However, the most intriguing observation from the comparison of EAS2 and OCP^O^ is an increased absorbance at 350 nm (black solid line, Fig 4b) of EAS2.

This 350-nm band yields the so-called ‘*cis*-peak’, a characteristic signature of *trans*-*cis* isomerization in carotenoids^36,37^. The IR spectrum of the corresponding EAS2 component in D_2_O buffer shows positive bands in the 1655-1641 cm^-1^ region while exhibiting a negative band at 1540-1550 cm^-1^(Fig 4c, red solid line). The positive band corresponds to the amide I signal. The position of the 1542-1550 cm^-1^ bleach is insensitive to D_2_O / H_2_O exchange (Extended Data Fig 4f, red solid line) and agrees with the frequency of C=C stretches in canthaxanthin ^38^ and other carotenoids^39^. In carotenoids, the oscillator strength of C=C stretches is known to be affected upon C=C reorientation or reorgainsation^40^. Therefore, the presence of the 1542-1550 cm^-1^ bleach in the EAS2 component could be a signature of the BP isomerization of CAN as seen in Fig 1c in I1. The EAS3 component confirms that BP isomerization is a transient phenomenon in 2.5 min leading to a ‘relaxed “ state where C9’-C8’ and C7’-C6’ single bonds are back in *trans* and *cis* conformations, respectively, as can be concluded from a decrease of the 1542 cm^-1^ bleach (162-s, Fig 4c, blue dotted line). An additional UV/Vis measurement with a shorter illumination period confirms that the back relaxation of the two concerted isomerization steps is thermally driven (Extended Data Fig 4 a, see Supplementary Information for the detailed explanation). To confirm the FTIR assignments, we also performed harmonic frequency calculations of CAN in the dark conformation (CAN_Dark_) and in the BP isomer (CAN_I1_) using the carotenoid coordinates from Fig 1c after optimisation using redundant coordinates (Fig 4d). The major frequency of the C=C stretches coupled with the C=O stretches in CAN_Dark_ is 1542 cm^-1^ (Fig 4 d, cyan sticks), while in the CAN_I1_ state, its oscillator strength decreases and its frequency redshifts to 1550 cm^-1^ (Fig 4 d, orange sticks). The resulting calculated difference signal (CAN_I1_-CAN_Dark_) is characterized by the 1542 cm^-1^ bleach (Fig 4d, red dotted line). Our spectroscopic results on OCP solution, supported by frequency calculations, thus confirm that BP single bond isomerization is a light-driven process occurring also in physiological conditions.

Here we report photokinetics of OCP with CAN, but the OCP has been shown to bind and be active with various keto-carotenoids (3′-hydroxyechinenone, 3’-hECN; echinenone, ECN; as well as CAN)^10,18,41^. The 350 nm peak observed here was previously reported in various carotenoids^36,37^, therefore it could be a BP isomerization signature in all OCP keto-carotenoid constructs. On the other hand, the position and oscillation strength of C=C stretch is carotenoid-specific: the 1542-1550 cm^-1^ bleach observed here is unique for CAN and e.g. C=C stretch in ECH is observed at lower amplitude and higher frequencies^42^. Therefore, the spectrum of BP conformation change in IR region is keto-carotenoid-specific.

The EAS2 component also contains positive amide-I FTIR signals peaking at 1651 cm^-1^ and 1641 cm^-1^ due to a bleaching signal emerging at later stages (EAS3, Fig 4c, blue dotted line). These signals, assigned to amide-I peaks based on previously reported IR signals of OCP WT and mutants^19^, are indicative of H-bond rearrangements which are seen in the I1 crystal structure (Extended Data Table 4). The positive amide-I FTIR signals can be assigned to the new polar contacts appearing in I1 (EAS2, 38 s, Fig 4c). Negative amide-I FTIR signals in the EAS3 component indicate the loss of these polar contacts as seen in I4 (Extended Data Table 4).

In line with previous studies^13,19,32^ the absorption peak at 1651 cm^-1^ is insensitive to D_2_O/H_2_O exchange, unlike that at 1645 cm^-1^ (Fig 4c and Extended Data Fig 4 g, Supplemental Information). We have therefore tentatively assigned the 1651 cm^-1^ and 1641 cm^-1^ (D_2_O)/1645 cm^-1^ (H_2_O) bands to a buried α-helix and to a solvent exposed α-helix of NTD respectively

In agreement with ^32^ we assign ∼1600−1645 cm^−1^ shoulder in FTIR (in H_2_O) to vibrational motions of HOH (Extended Data Fig 4f). See Supplementary Information for more details regarding the contribution of water molecules to FTIR data. In support of this assignment in Extended Data Fig 4f, we observed that the presence of the 1600-1640 cm^-1^ band in the H_2_O data (Extended Data Fig 4 f-EAS2 and EAS3) correlates with the light-driven displacement of water molecules observed in the crystallographic data on the same time scale (Extended Data Fig 5, Extended Data Fig 6-I2 and I3). The rearrangement of water molecules affects the two water clusters CL1 and CL2 each centred on HOH46 and HOH152 respectively. The strongest DED signals are observed in CL1, which is located on the interface between the two domains (Extended Data Fig 5).

Contrary to a previous suggestion^32^, our data does not support involvement of the C-terminal tail to changes of the 1645 cm^-1^ peak, since no structural rearrangements are observed in this region as can be judged from DED maps and DED amplitudes in the crystallographic data corresponding to the same time delays (I1-I2, Fig 2, Extended Data Fig 2 b). In line with the final I4 crystal structure, where the CTD translocates away from the NTD without CTD rearrangement, we do not observe CTD characteristic ß-sheet-related FTIR signals in the 1640-1600 cm^-1^ region (Fig 4c, violet dashed line). Since as was shown previously^11,15^ the process of domain separation occurs without substantial unfolding of the secondary structure, we observe very low amplitude of the FTIR signals in the amide I region (1640-1660 cm^-1^). The EAS4 (infinity/steady-state), therefore, represents the CTD and NTD complete dissociation^11^, which is not observable in the crystallographic data due to the crystal packing constraints ^18^. What is more, the bleach around 1550-1575 cm^-1^ of EAS4 (Fig 4c, violet dashed line) is indicative of some isomerization process. It is in agreement with a previous study in which C6−C7 s-*cis* CAN was observed in RCP^15^. RCP is a truncated OCP fragment which is assumed to be a native OCP^R^ state after domain separation^15^. To confirm our 1550-1575 cm^-1^ bleach assignment, we also performed Harmonic frequency calculations (HFC) at the B3LYP/6-311+G level of IR spectra for CAN carotenoid in the RCP conformation (Extended Data Fig 4 h-j). The results demonstrate that the FTIR difference signals between CAN_RCP_ and CAN_Dark_ would result in the 1550-75 cm^-1^ bleach which clearly has a more pronounced shoulder in longer wavenumbers (violet dotted line, Extended Data Fig 4 j) as compared to the 1542 cm^-1^ bleach characteristic of BP isomerization as shown on Fig 1 c. Our HFC predictions for the CAN_RCP_ -CAN_Dark_ difference IR spectrum therefore agrees very well with the EAS4 assignments.

Except for the final step of CTD from NTD separation, which is not achievable in crystallography due to crystal packing constraints, amide-I FTIR kinetics data and assignments agree well with our crystallographic observations, indicating that the same photoinduced structural rearrangements occur both in OCP solutions and in crystals. This work provides the structural basis for the initial reaction in OCP light-activation. Our X-ray data demonstrates that the initial photoproduct is the s-isomerization along C9’-C8’ and C7’-C6’ single bonds of CAN carotenoid in a BP manner. To propose how these reorientation motions could be possibly driven by a photon absorption, the carotenoid electronic structure should be considered. Light absorption in carotenoids like in other linear polyenes, is accosiated with S_0_(1^1^Ag^-^) → S_2_(1^1^Bu+) transition, while the optically forbidden S_1_ dark state (2^1^Ag^−^) is rapidally populated through internal conversion. Multiple reports indicate that the conical intersection driving the S_1_(2^1^Ag^−^) → S_0_(1^1^Ag^−^) trasition in polyenes of various length corresponds to a -(CH_3_)-kink along the conjugated backbone where several sequential bonds are totally or partially twisted^43,44^. We therefore put forward a hypothesis that the constrained protein environment of OCP drives the generation of the BP photoproduct as observed in I1 intermediate via CAN S_1_ state.

UV/Vis and IR spectroscopic studies in combination with harmonic frequency calculations confirm the reaction mechanism in crystals and solution. In response to the light-induced isomerization, structural changes propagate from the NTD to the CTD. The presented evidence shows that the H-bond rupture between the carotenoid and the CTD protein moiety does not initiate light activation but is a secondary event which follows post-isomerization NTD rearrangement.

While OCP has been shown to be active with different keto-carotenoids resulting in identical photocycle intermediates^19,32^, OCP with 3’-hECN was found to be the dominant form of OCP in native cyanobacteria^45^. Since the primary OCP photoproduct (CAN_I1_) demonstrates two single bonds rotation around C9’ and C6’ carbons (Fig 1 c) in NTD during BP isomerization and does not involve β2 ring, the mechanism observed here should be universal for OCP with different keto-carotenoids and, therefore, naturally occurring in cyanobacteria during photoprotection.

Previously BP photoisomerization was observed in another conjugated chromophore, retinal^26^, which is synthesized from a carotenoid by irreversible oxidative cleavage^46^. However, there the process involves two-double-bond isomerization. The mechanism observed here involves two orchestrated single-bond isomerization steps in a BP manner and has never been reported before. Therefore, this finding is of a great importance for our understanding of carotenoid photophysics and evolution of photosensing of conjugated systems, in general. Furthermore, the discovery of the overall OCP activation mechanism provides for the first time the foundation for gaining control and manipulation of the light regulation function of OCP with potential applications in optogenetics, bioimaging and agriculture.

## Methods

### OCP expression

E. coli BL21-Gold (DE3) cells from Agilent Technologies were used for OCP expression. The OCP-encoding gene slr1963 from Synechocystis PCC 6803 was inserted between the NdeI/XhoI sites in the pET28 vector resulting in OCP-pET28 plasmid carrying G-S-S 6X His Tag-OCP at the NTD. To promote canthaxanthin (CAN) incorporation in vivo, the OCP-pET28 plasmid was co-expressed with pAC-CANTHipi plasmid (gift from Francis X Cunningham Jr, Addgene plasmid # 53301; http://n2t.net/addgene:53301; RRID:Addgene_53301). The transformed cells were then grown in the presence of kanamycin (25 μM/ml) and chloramphenicol (25 μM/ mL). Induction was carried out in LB medium at 37 °C for 3-4 hours until the optical density at 600 nm (OD_600_) of 0.6-0.8 was reached. Then 0.2 mM IPTG was added to trigger OCP expression and the culture was grown for 16-18 hours at 26 °C before harvesting the next day. After IPTG addition, E.coli cells were grown in the dark to prevent any OCP^O^-OCP^R^ conversion. Cells were harvested and pelleted by centrifugation for 15 minutes at 5,000 x g at 4°C and were subsequently stored at -20 °C.

### OCP purification

Cell pellets were thawed and re-suspended in lysis buffer (40 mM Tris pH 8, 300 mM NaCl and 10 % glycerol) supplemented with DNase I (0.1 mg/ml) (Roche) and protease inhibitors (1 tablet per 100ml) (Sigma) prior to cell disruption. All purification steps were carried out at 4 °C. Cell lysis was performed with two passes in a cell disrupter (Constant Systems Ltd.) at 40 kpsi. Broken cells then underwent ultra-centrifugation at 142,000 x g for 60 minutes before collecting the cell free extract and incubating it with pre-equilibrated HisTrap Ni^2+^ resin (GE Healthcare) in a 3:1 volume ratio for 2 hours under gentle shaking. Once the slurry was poured in a glass column, His-tagged OCP was washed with five column volumes (CV) of lysis buffer (wash 1) followed by three CV of lysis buffer supplement with 20mM imidazole (wash 2). The protein was washed off with elution buffer (40 mM Tris pH 8, 300 mM NaCl, 10 % glycerol, and 200 mM imidazole). Holo-OCP was subsequently separated from apo-OCP using anion exchange chromatography (DEAE 650 M) (Tosoh, Japan) by performing a 0-100 mM NaCl linear gradient. The concertation of Holo-OCP was determined using the maximum extinction coefficient of CAN (118,000 M^-1^·cm^−1^)^48,49^ and the known 1:1 binding stoichiometry of CAN:OCP.

### Protein crystallization

OCP was washed with washing buffer (40 mM Tris pH 8, 10 % glycerol) prior to crystallization using the sitting drop vapor diffusion method based on the protocol reported in^15^ and described below: 2 μL of OCP at 3 mg/mL was mixed with 1 μL of reservoir solution (100 mM sodium acetate pH 4.5, 10 % poly-ethylene glycol 20,000, 3 % glycerol) and the drops were left to equilibrate at 20 °C for 5 days. Cubic crystals were flash frozen in liquid nitrogen after being transferred to a cryoprotectant solution consisting of the crystallization solution with 30% glycerol.

### Absorbance measurements

#### Crystal pancakes

Slurries of OCP crystals were pelleted at 1,000 x g and 4°C for 2 minutes and the supernatant removed. Pellets were scooped out and squeezed between two siliconized microscope coverslips using a micrometre to apply controlled force before sealing the edges with superglue. This process ensured the production of a pancake of crystalline matter with a reasonably uniform OD at 410 nm. The final OD_410_ was typically around 0.2-0.3. Illumination was provided by a 10W 410nm Purple COB LED generating 100mW of 410 nm light at the interaction area. The same LED with similar power was used for illumination of single crystals before flash freezing.

#### Solutions

UV-VIS measurements were collected using an Agilent 8453 spectrophotometer. Samples with an OD_500_ = 0.4 were used in a 1 cm light path quartz cuvette. Samples were kept at 20 °C and illuminated with a 405 nm laser of 100mW (Fig 4 a-b and Extended Data Fig 4a) or variable power (Extended Data Fig 4b-c) for 10 min (Fig 4 a-b) or for 40 s (Extended Data Fig 4a). To confirm that the bicycle-pedal isomer relaxation is thermally-driven, the time-dependent UV-VIS spectra in Extended Data Fig 4 a were followed by 5 min of darkness after 40 s of illumination. Absorbance changes were collected at 0.5 s intervals.

#### FTIR measurements

Fourier-transform infrared (FTIR) spectra were recorded at a 1 cm^−1^ resolution using a Bio-Rad FTS 175C FT-IR spectrophotometer equipped with a mercury cadmium telluride (MCT) detector. Before measurements the instrument was purged with CO_2_-free dry air for 120 min. Absorption spectra were collected in the region between 1700 and 1500 cm^−1^. For FTIR measurements, OCP in its original buffer (40 mM Tris pH 8, 10 % glycerol) was exchanged into either H_2_O (40mM Tris pH 8) or D_2_O (40mM Tris pD 8) based buffers. 10 μL of sample previously concentrated to the maximum achievable concentration was placed into the Harrick cell fitted with a 6 μm spacer to obtain an OD_500_ around 0.2. First, FTIR was measured in a dark-adapted (non-illuminated) state consisting of 2000 interferograms. The sample was then illuminated at 405 nm with a 100mW laser. Each FTIR spectrum containing 100 interferograms was recorded in 35 seconds. A total of 16 spectra were collected sequentially for about 10 min after the onset of illumination.

### Global fit of spectroscopic data

The obtained absorbance changes were analysed globally using Glotaran software^33^ and the methodology described in ^34^. During the global analysis UV-VIS and FTIR datasets (both D_2_O and H_2_O) were linked together. The unbranched and unidirectional, so-called sequential model (EAS1→EAS2→… →EASn_comp_) was used, where EAS stands for Evolution Associated Spectra. During the global analysis, UV-VIS and FTIR datasets in both D_2_O and H_2_O buffers were linked together.

### Sample optimization and X-ray crystallography

OCP crystals had cubic morphology. The cubes with the length of not more than 30 μm were selected for the illumination experiment to allow light penetration through the crystal. Taking into account unit cell dimensions and CAN extinction coefficient (118,000 M^-1^·cm^−1^)^48,49^ OD of single crystals is ∼1.0. Therefore only a few percent of absorbed light are transmitted. As a result, overall populations of illuminated states do not exceed 30% in single crystals. Illumination was performed at RT using a 10W 410nm Purple COB LED generating 100mW of 410 nm light at the interaction area. Great care was taken so that the crystal was evenly exposed to the set illuminated intensity over the desired illumination time. After preillumination the crystals were flash frozen in liquid nitrogen after being gradually transferred to a cryoprotectant solution (crystallization solution with 30% glycerol). To be sure that no build-up intermediates would relax the crystals were also illuminated while moving into cryoprotectant or in liquid N2. The crystals were illuminated for different final time illuminated periods: 1 min minute (I1), 2 minutes (I2), 5 minutes (I3) and 10 minutes (I4). After the illuminated crystals have been cryotrapped they were sent to Diamond Light source for data collection. All X-ray diffraction data collection was carried out at i03 and i04 beamlines (Diamond Light Source, UK).

### X-ray data refinement

OCP crystals belonged to the P3_2_21 space group with a = 82.5 Å, b = 82.5 Å, and c = 87.3 Å. Diffraction data were integrated through xia2 pipeline with DIALS^50^. The structures were solved by molecular replacement with MOLREP ^51^ using a starting model based on the PDB ID 4XB5. Refinement was performed with REFMAC^52^ alternating with / model building using 2Fo-Fc and Fo-Fc maps visualized in COOT^53^. The final model contains one OCP molecule in the asymmetric unit. Statistics for diffraction data collection, structure determination and refinement are summarized in Supplementary Information.

Difference electron density (DED) maps, calculated for various dark datasets (F_Dark_–F_Dark_) reveal no map features, indicating that the different “dark” structures are identical and do not exhibit any spatial arrangement differences as expected (Extended Data Fig 2 a). DED maps were generated using Q-weighted method (see Extrapolated maps and coordinates). In all four cases, the obtained difference maps in different states are highly reproducible (Extended Data Fig 2). The results indicate that the signals must be generated by actual biological phenomenon during OCP light-activation and do not arise from statistical noise. Similar difference signals were also obtained using phenix.fobs_minus_fobs_map method. Figures of crystal structures and difference maps were visualized using PyMOL Molecular Graphics System (Version 2.5.0, www.pymol.org).

Difference density amplitudes were obtained according to ^47^. In short, difference amplitudes A_j_(a_i_) and difference Fourier electron density maps within spheres of radius 2.0 Å centred upon atom a_i_ were averaged using a 3 σ cut-off and a 0.5 Å grid spacing.

### Extrapolated maps and coordinates

To generate extrapolated maps firstly, DED were calculated using the Fourier transform of weighted difference structure amplitudes based on the Bayesian statistics analysis by Ursby *et. al* ^27,54^. For each corresponding structure factor difference (ΔF) between the light (F_L_) and dark (F_D_) data, a weighted factor (w) was calculated and applied to the difference as implemented in ^55–57^. Light, dark and calculated structure factors were combined using CAD and scaled using the anisomorphous option in ScaleIt ^54^. This placed each set of structure factors onto a normalised absolute scale compared to the calculated structure factors (F_c_ calculated using SFALL). The final real-space map was calculated using FFT. Phases from the dark state structure, corresponding to unilluminated OCP crystals were used.^54^ To identify small structural changes in the illuminated data, extrapolated electron density maps were generated and extrapolated coordinates then refined to these maps. Difference structure factors were added in a linear combination in multiples of N_EXT_ to the calculated dark structure (F_Dark_) in order to generate extrapolated structure factors (F_EXT_). The negative electron density around areas of interest in the protein were then integrated. The point where negative density started to build-up was used to determine a characteristic N_EXT_ value which was then used to generate the final extrapolated structure factors. The population transfer (PT) was around 15-30% as can be approximated from the N_EXT_ value (PT = 200/ N_EXT_)^57^. Coordinates were refined against the extrapolated structure factors using five cycles of rigid body refinement in REFMAC^58^. The real space refinement tool on COOT^53^was used to fit the carotenoid and neighbouring amino acids to the extrapolated electron density. These coordinates were then saved and used for the occupancy refinement against the ground state/dark structure for I1-I4 datasets. Occupancy refers to the proportion of extrapolated coordinates (light, I1-I4) compared to ground state coordinates (OCP^O^). Both R and R_free_ minimise at the occupancies: Dark=0.84/Light=0.16 (I1), Dark=0.78/Light=0.22 (I1), Dark=0.75/Light=0.25 (I1) and Dark=0.68/Light=0.32 (I4).

### Phasing experiment

The crystals were harvested using LithoLoop (Molecular Dimensions) sample mounts on specialised I23 copper sample assemblies. Data were collected on the long-wavelength beamline I23^59^ at Diamond Light Source with X-ray energies of 2.9 keV (λ = 4.28 Å) and 2.8 keV (λ = 4.43 Å) using a semi-cylindrical PILATUS 12M detector (Dectris, CH). These energies were selected as they are above and below the Cl absorption edge (E = 2.82 keV/ λ = 4.39 Å) respectively, thus allowing for Cl identification during analysis of anomalous difference Fourier maps. Sweeps of 360° with an exposure time of 0.1s per 0.1° image were taken using the same crystal. To ensure completeness, two datasets for each energy were collected with the multi-axis goniometer at angles κ=0° and φ=0° and, subsequently, κ=-30° and φ=60°. These datasets were interleaved to mitigate radiation damage effects.

For data processing, data integration was performed with XDS and XSCALE programmes. The origin of the PDB model was corrected using POINTLESS, AIMLESS and MOLREP. Phased anomalous difference Fourier maps were produced using ANODE. A cut-off of 6.0 σ was used to identify sites of anomalous contribution.

### Harmonic frequency calculations

The Gaussian 06 software package^60,61^ and the Becke-3-Lee-Yang-Parr functional ^62^ were used with the 6-311+ G basis set. Theoretical IR values for frequencies and intensities in CAN were calculated for the three states CAN_Dark_ (Fig 4d), CAN_I1_ (Fig 4d), CAN_I4_ (Extended Data Fig 4 i) and for the RCP structure as reported in ^15^ (Extended Data Fig 4 j). FTIR spectra were derived from the theoretical IR values using Gaussian program by assigning FWHM withspreading the intensity over a Lorentzian function. The Lorentzian line width for each absorption peak in the calculation was taken to be 4 cm^− 1^. The optimization of molecular geometry was performed with fixing all CAN dihedral angles. As a control, FTIR spectra were also calculated for CAN_I1_ and CAN_Dark_ optimised geometries where dihedral angles were not fixed (Extended Data Fig 4 h). The CAN_I1_-CAN_Dark_ difference signal in Extended Data Fig 4 h results in a similar bleach observed in Fig 4d but with lower amplitude and a shift to lower wavenumbers.

A scaling factor of 0.97^63–65^ was used to convert the calculated theoretical frequencies to FTIR spectra.

## Acknowledgements

The data were collected remotely at Diamond Light Source (BAG proposal number mx17221). This project was supported by the EMBO long-term fellowship (EMBO ALTF 244-2017) and has received funding from the European Union Horizon 2020 research and innovation programme under the Marie Sklodowska-Curie grant (agreement No. 839389) to V.C. This work was supported by the Leverhulme Trust award RPG-2018-372. The crystallisation facility at Imperial College was funded by the BBSRC (BB/D524840/1) and Wellcome Trust (202926/Z/16/Z). We thank Violeta Cordon Preciado for technical assistance.

## Author contributions

V.U.C. designed the project. V.U.C. expressed and purified OCP protein, prepared OCP crystals, collected and processed X-ray data and UV-VIS/FTIR data. J.M.B assisted with creating Q-weighted maps and extrapolated coordinates. A.F. and J.J.vT. performed Harmonic Oscillator Calculations. R.M.M. assisted with cryo trapping experiment and X-ray data processing. M.A.P. assisted with the designing of OCP expression system in E.coli, while K.M. provided advice and support in obtaining OCP crystals. C.O. and A.W. performed long wavelength experiments. V.U.C. wrote the manuscript with the input from J.B., R.M.M., A.F., C.O. and A.W. J.M.B., R.M.M., A.F., C.O., A.W. and in particular K.M., V.U.C. and J.J.vT. contributed to the review and editing. J.J.vT. supervised the project.

## Data and materials availability

Coordinates and structure factors have been deposited in the Protein Data Bank under accession code 7ZSF (OCP^O^), 7ZSG(I1), 7ZSH(I2), 7ZSI(I3) and 7ZSJ(I4). Extrapolated maps, extrapolated coordinates and other data associated with this manuscript are available from the authors on reasonable request.

## Supplementary Information

Supplementary Information is available for this paper.

## Extended Data

**Extended Data Figure 1.**
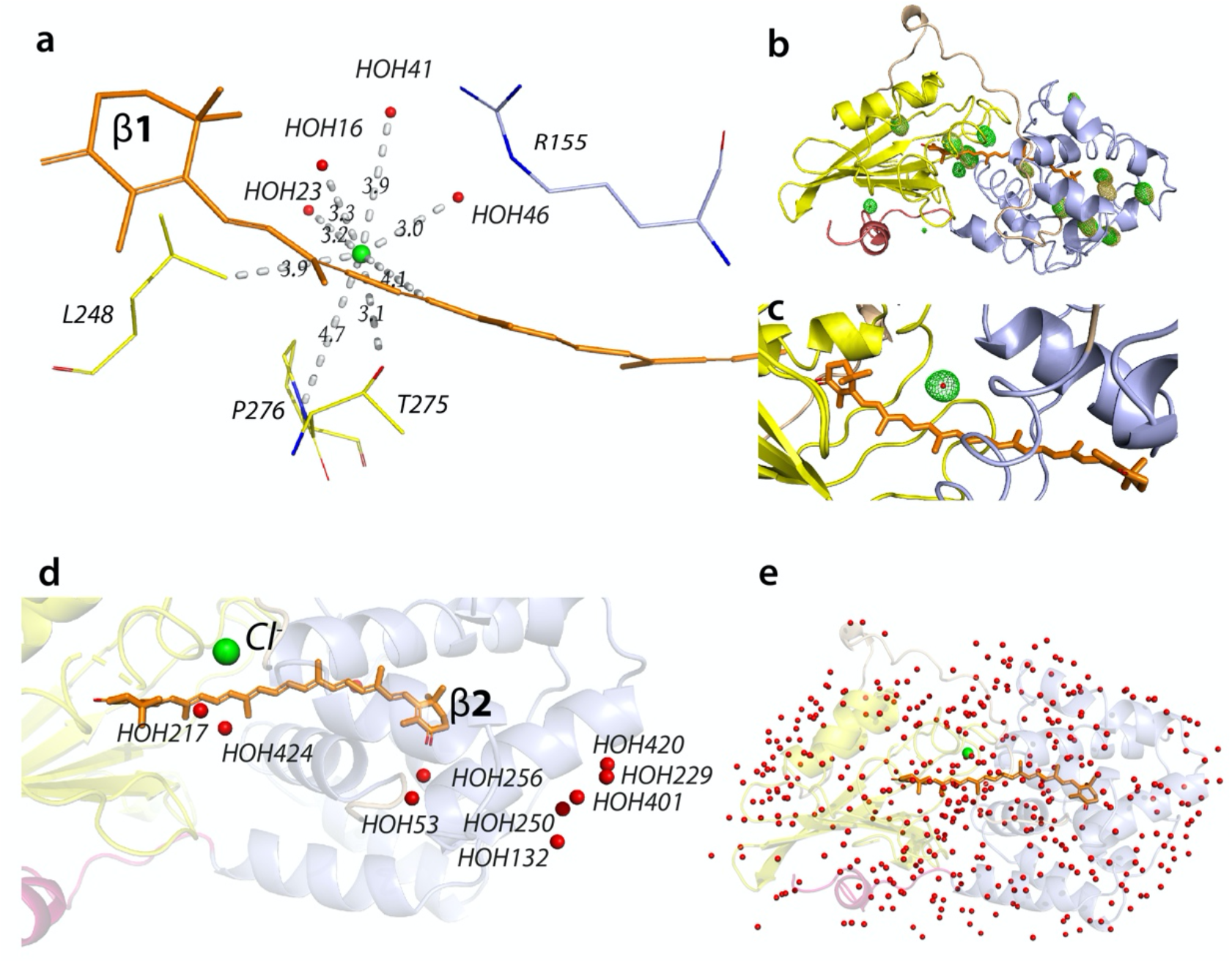
Identifying Cl– location. (a) Cl-(green sphere) is resolved in the CTD using Long-wavelength experiment for Cl-identification. Cl-coordinates four water molecules (red spheres) through ion-dipole interactions at distances of 3.0-3.2 Å (HOH16, HOH23, HOH46) and 4.0 Å (HOH41). (b) Overall structure anomalous difference density Fourier maps at 2.9 keV (green) and 2.8 keV (light orange). The regions where green density is present and light orange density where it is absent indicate the presence of Cl (red dot). In the instances where the Cl atom could not be modelled due to a sigma greater than 6, dummy atoms have been placed to indicate the potential location (dots). (c) Close-up view of the modelled Cl-site (red dot) with the 2.9 keV anomalous difference Fourier map density surrounding it (green). (d) The newly identified water molecules as described in Extended Table 2. (e) A total of 401 water molecules (red spheres) were resolved in the OCP structure reported herein.

**Extended Data Figure 2.**
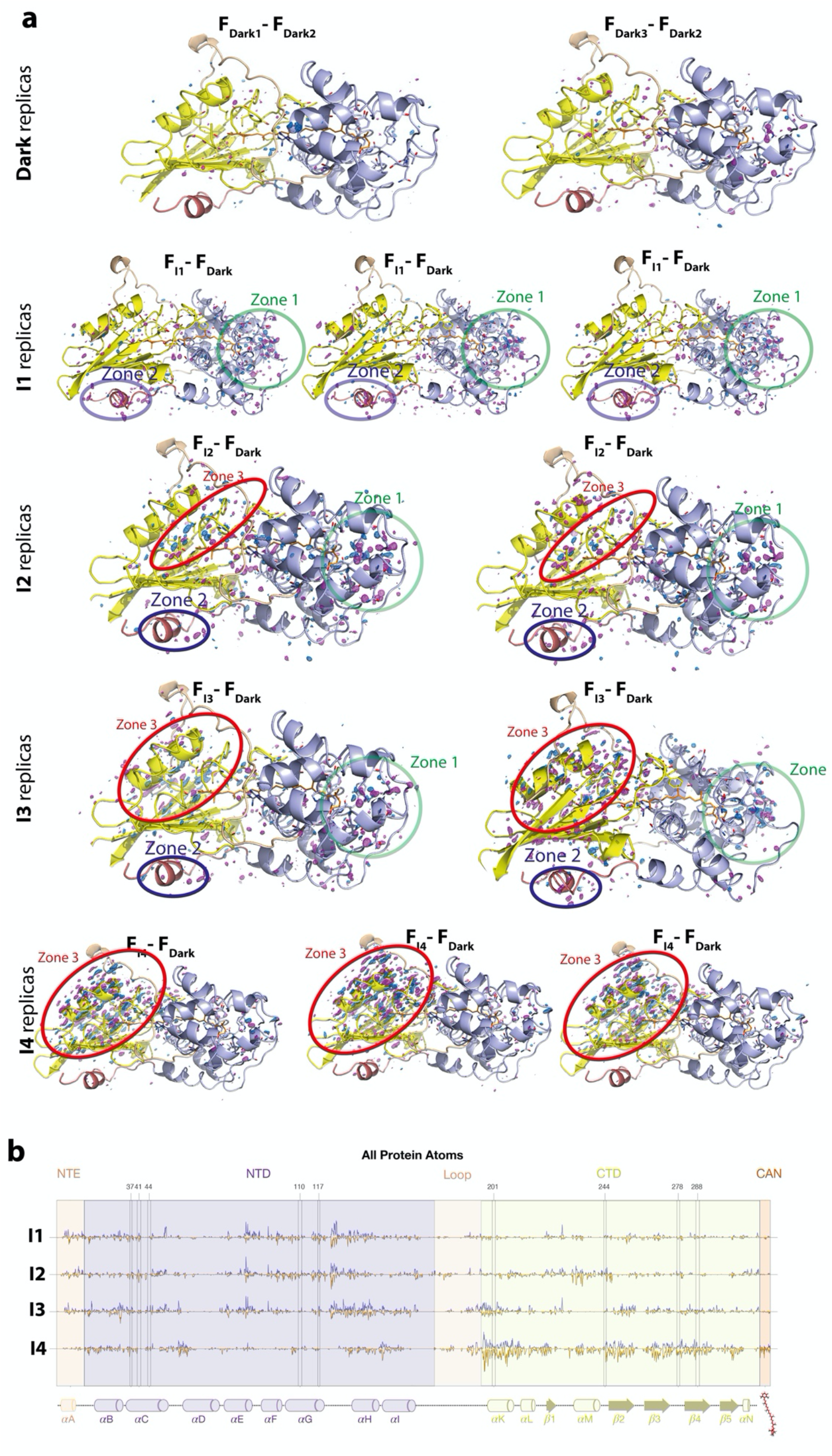
(a) DED maps contoured at ±3.5σ (blue and magenta correspond to the appearance and disappearance of density, contoured at 3.5σ). Dark state DED signals contoured at (±)3.5σ. The F_Dark1_–F_Dark2_ and F_Dark3_– F_Dark2_ exhibit marginal DED signals. The F_I1_– F_Dark_, F_I2_–F_Dark_, F_I3_–F_Dark_ and F_I4_–F_Dark_ datasets show reproducible strong DED signals in the three different zones. The following domains undergoing photoinduced conformational changes are represented in oval shapes: Zones 1 (NTD), Zones 2 (NTE) and Zones 3 (Alpha-helices in CTD) (b) Difference electron density amplitudes calculated as described in the X-ray data refinement section of the Materials and Methods.

**Extended Data Figure 3.**
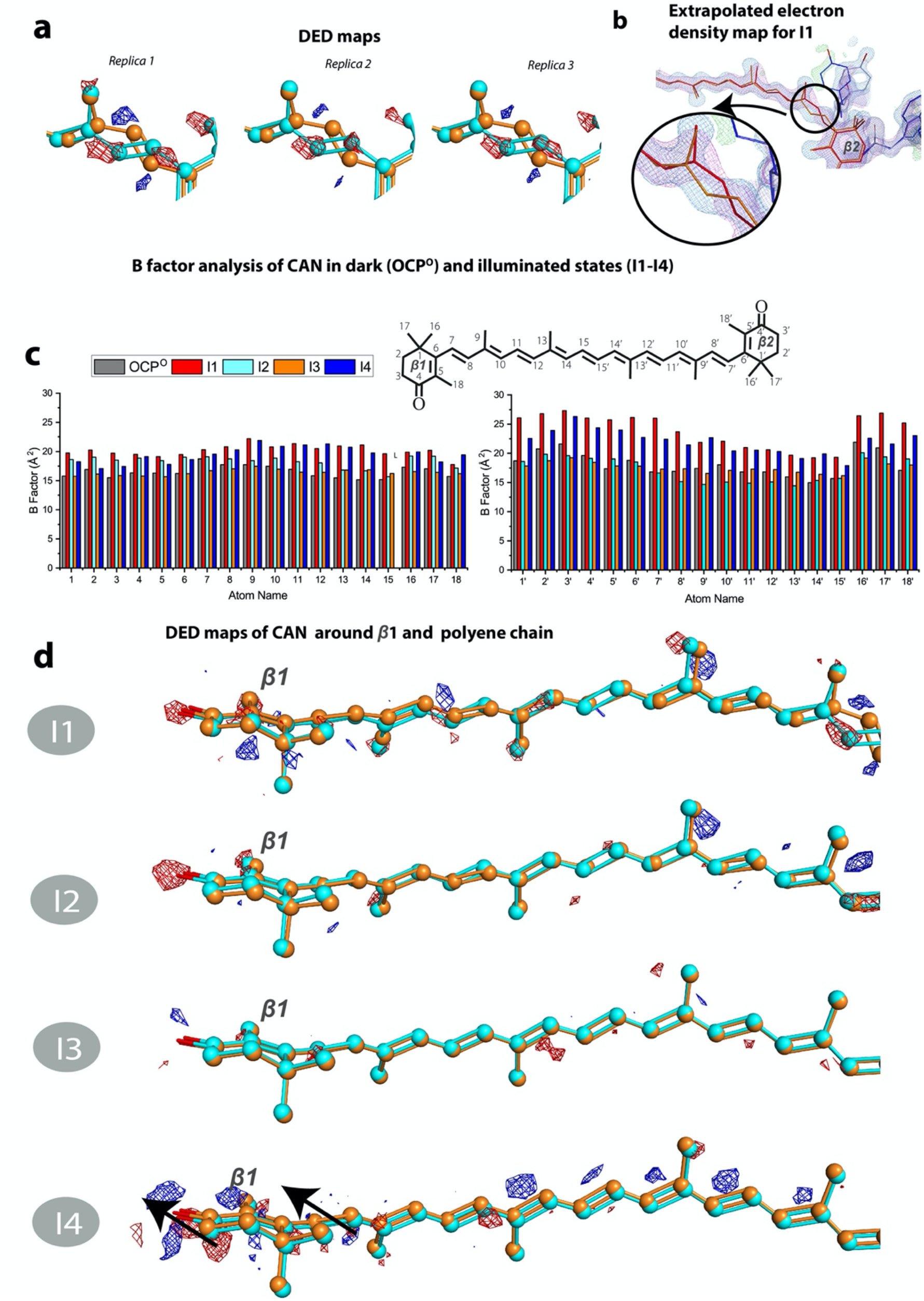
(a) Reproducibility of isomerization signal. Similar DED signals assigned to isomerization were observed for three different I1 datasets. (b) Extrapolated maps and coordinates of CAN. (c) Atomic B factors for CAN comparing the OCP° (dark) structure to the photoproduct states in I1, I2, I3 and I4 (d) DED signals around β1-ring and polyene chain of CAN in I1-I4 states.

**Extended Data Figure 4.**
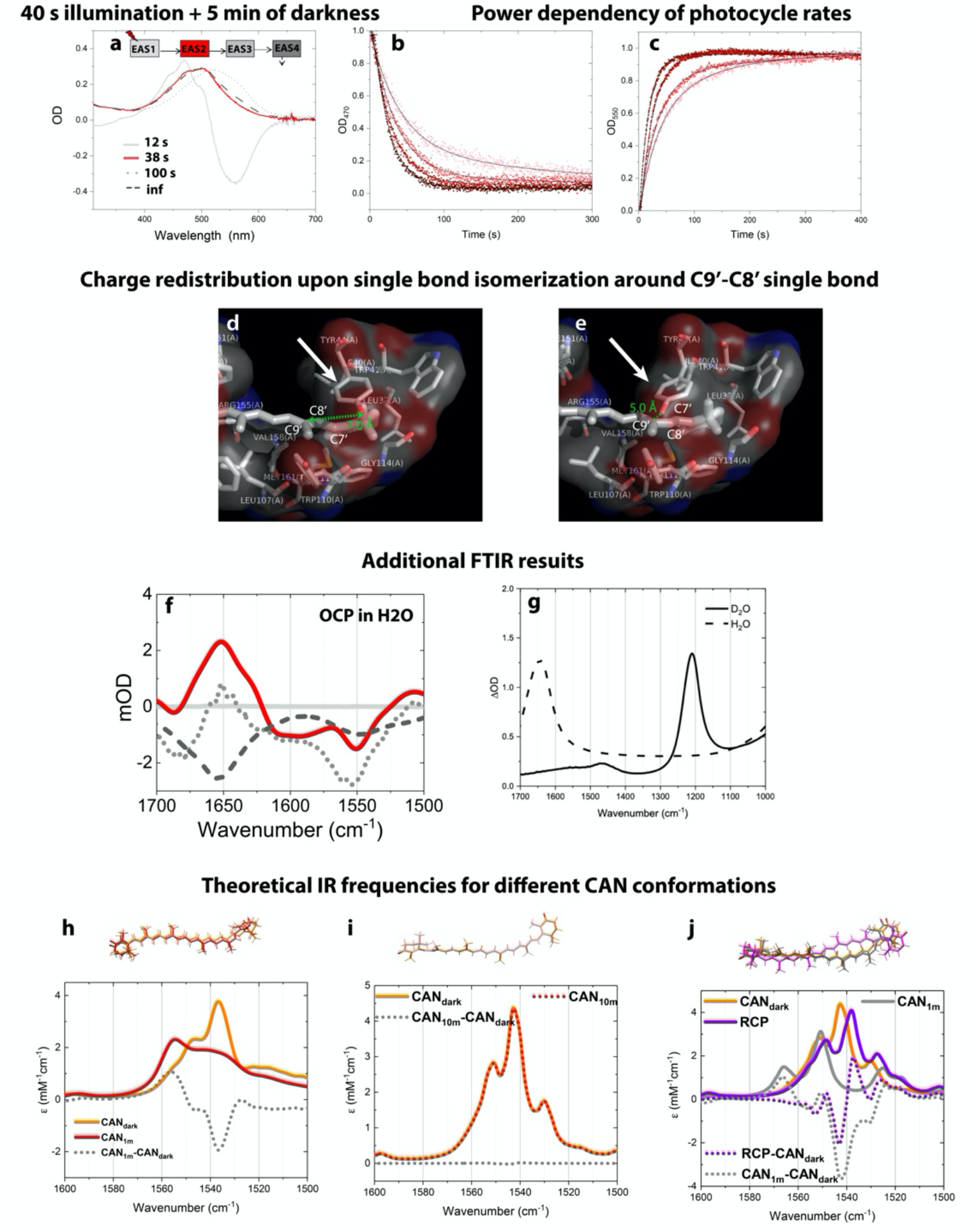
(a) Results of the global fit of UV-VIS absorption kinetics in solution upon 40 s of illumination followed by 5 min of darkness. The kinetic model used for the fit is indicated at the top of the graph which also shows the lifetimes. (b and c) Power dependency profile of photocycle intermediate accumulation rates upon continuous illumination with violet light (410 nm) as monitored by 470 nm bleach (b) and 550 nm rise (c). Power densities used going from dark red to pink: 3.5 mW cm^-2^, 2.8 mW cm^-2^, 1.0 mW cm^-2^ and 0.5 mW cm^-2^. Black lines represent double exponential fits of the curves. (d-e) Zoomed view of the carotenoid tunnel near the β2 ring in OCP^O^ (d) and I1 (e) states. The two single bonds isomerization (C9’-C8’ and C7’-C6’) in a bicycle pedal manner causes charge redistribution in the C9’-C8’=C7’-C6’ CAN moiety. (f) Global analysis results obtained for OCP FTIR signals (H_2_O) from the simultaneous global fit of UV-VIS/FTIR data. (g) FTIR spectra of H_2_O (dashed line) and D_2_O (solid line) alone. (h-j)Theoretical IR intensities were calculated using Harmonic frequency calculations (HFC) at the B3LYP/6-311+G level. (h) IR values were calculated for CAN in the two states CAN_Dark_ and CAN_I1_ when dihedral angles were freed in the calculation of the optimised geometry. (i) Calculated IR spectra for CAN_dark_ and CAN_10m_. (j) Calculated IR spectra for CAN_dark_, CAN_1m_ and RCP.

**Extended Data Figure 5.**
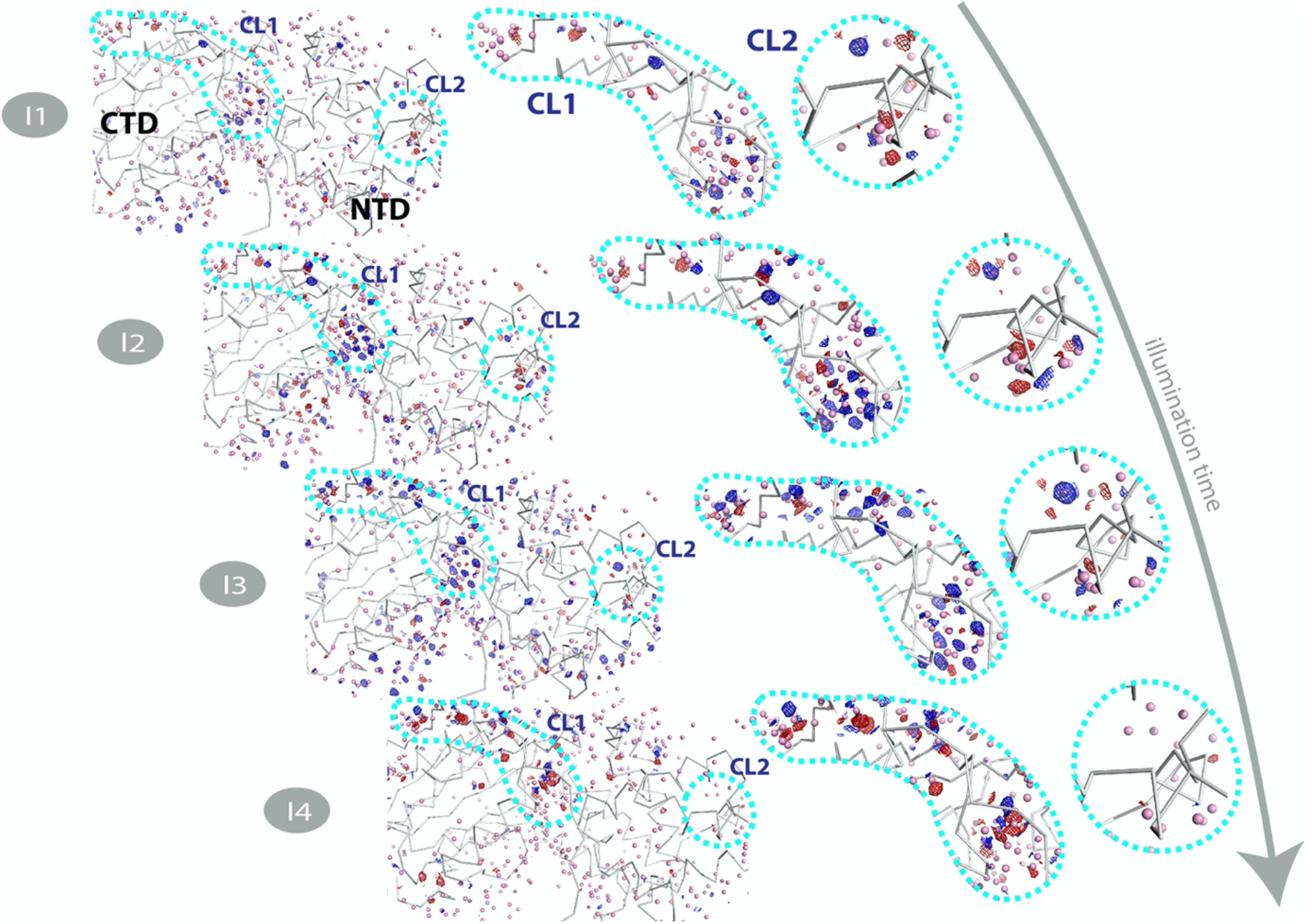
Light-driven water molecules redistribution in the OCP crystal structures. Q-weighted DED maps associated with the four pairs I1, I2, I3 and I4 (in blue and red, contoured at ±3.0σ). Amino acids are represented by grey ribbons, while water molecules are shown as pink spheres respectively. CL1 and CL2 indicate water clusters.

**Extended Data Figure 6.**
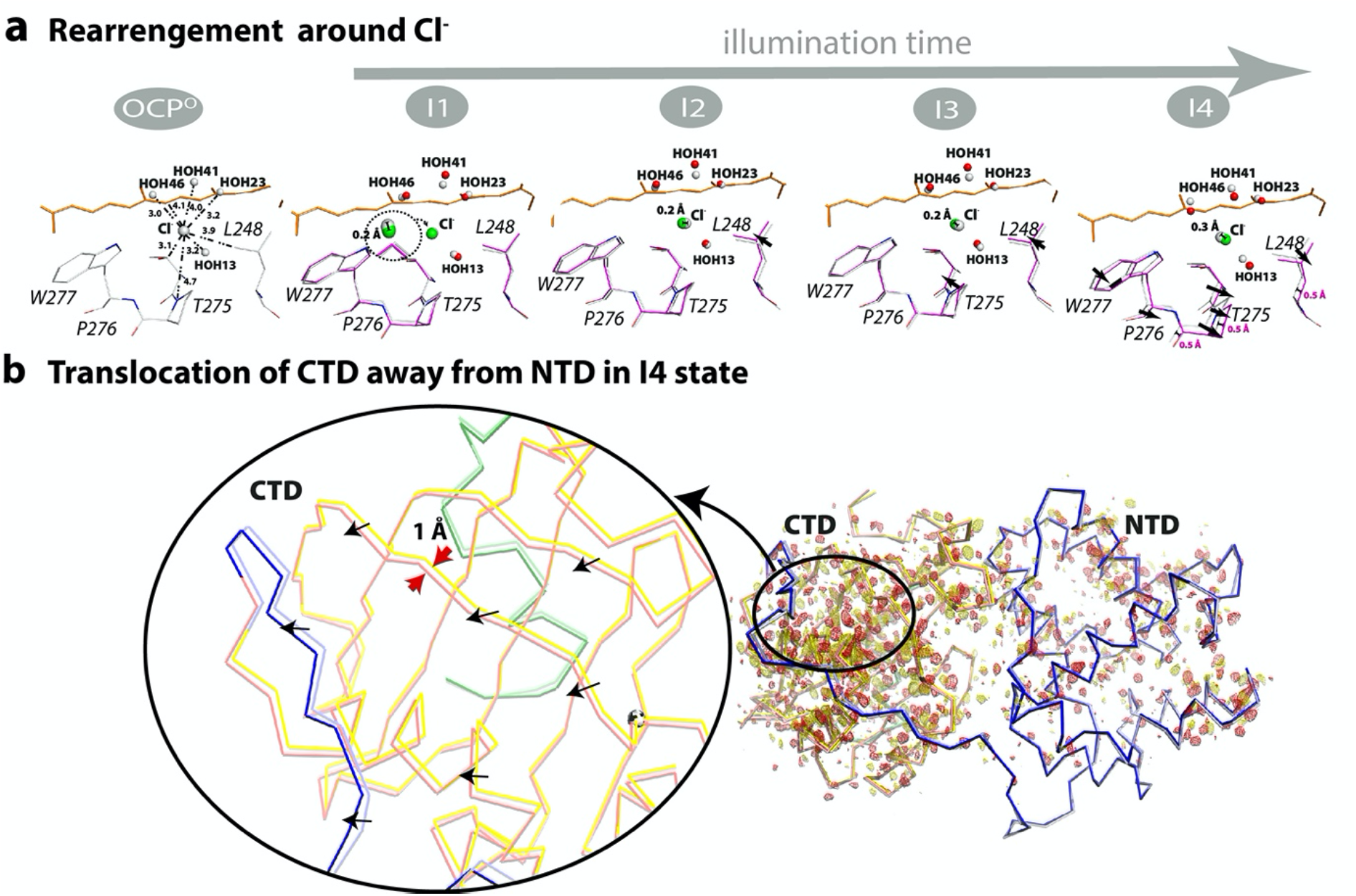
(a) Light-driven structural changes in I1-I4 states. Features shown are within a 4 Å radius from the Cl^-^ ion. The blue and red mesh represent DED maps contoured at ±3.0σ. Dark state amino acids, water and Cl^-^ coordinates are shown in grey while light state extrapolated coordinates are shown in pink sticks, red (HOH) and green (Cl^-^) spheres, respectively. (b) Translocation of CTD from NTD in I4. DED maps obtained for F_I4_–F_Dark_ (10 min) contoured at ±3.0σ (yellow and red for the appearance and disappearance of density respectively) and extrapolated coordinates for I4 vs OCP^O^ showing a 1 Å translocation of CTD away from NTD.

**Extended Data Table 1:** Comparison of canthaxanthin dihedral angles observed in OCP structures in the three states: Dark, I1 (1 min of illumination) and I4 (10 min of illumination). Dihedral angles were measured using atomic PDB coordinates for carbon atoms. Deposited dihedral angles for CAN-binding (PDB entry 4XB5), ECN-binding (3MG1) and 3’-hECN-binding (1M98) OCPs are also included for comparison.

|  | Dark (this study) | 1m (this study) | 10m (this study) | Syn OCP<sub>CAN</sub> (4XB5) | Syn OCP<sub>ECN</sub> (5TUX) | Syn OCP<sub>ECN</sub> (3MG1 - CA) | Syn OCP<sub>ECN</sub> (3MG1 - CB) | Arth OCP<sub>hECN</sub> (5UI2 - CA) | Arth OCP<sub>hECN</sub> (5UI2 - CB) |
| --- | --- | --- | --- | --- | --- | --- | --- | --- | --- |
| C5-C6-C7-C8 (β1 ring) | 153 | 157.8 | -35.6 | 116.3 | 131.8 | 146.4 | 148.6 | 118.5 | 131.9 |
| C6-C7-C8-C9 | 160 | 163.1 | 159.9 | 175.4 | 176.7 | 171 | 172.2 | 172.8 | 174.7 |
| C7-C8-C9-C10 | 167 | 160.4 | 167 | -155.3 | -168.7 | -179.5 | 179.5 | -163.7 | -176.8 |
| C8-C9-C10-C11 | 164.9 | 162.4 | 164.9 | 175.8 | 174.9 | 169.3 | 170.9 | 177.9 | 171.1 |
| C9-C10-C11-C12 | -176.1 | -166.1 | -176.2 | 178.8 | -179.5 | 173.2 | 173.5 | 165.2 | 173.6 |
| C10-C11-C12-C13 | 176.9 | -172.8 | 177.0 | 174.4 | 173.7 | 169.6 | 171.4 | 175.4 | 175.5 |
| C11-C12-C13-C14 | 177.6 | 163 | 177.7 | 167.8 | 170 | 172.8 | 178 | 177.5 | 175.9 |
| C12-C13-C14-C15 | 172.7 | 175.4 | 172.7 | 173 | 172.8 | -177.3 | -178.8 | 171 | 170 |
| C13-C14-C15-C15' | 175 | -176.2 | 175 | -171.4 | 177.8 | -175.2 | 179.8 | -175.7 | 171.5 |
| C14-C15-C15'-C14' | 172.4 | 144.9 | 172.4 | 172.7 | 171.8 | 168.7 | 167.7 | 170.7 | 171.4 |
| C15-C15'-C14'-C13' | -175.2 | 163.8 | -175.2 | 169.8 | -178.7 | 178.9 | 178 | 179.6 | -168.2 |
| C15'-C14'-C13'-C12' | 171.5 | -177.5 | 171.5 | 173 | 172.2 | 170.9 | 170.4 | 172.8 | 169.8 |
| C14'-C13'-C12'-C11' | -170.9 | -148.1 | -170.9 | -153.3 | -165.9 | -172.4 | -171.5 | -161.6 | 171.4 |
| C13'-C12'-C11'-C10' | 172 | 171 | 172 | 173.1 | 172.8 | 171.4 | 171 | 174.8 | 170.9 |
| C12'-C11'-C10'-C9' | -175.9 | 167.5 | -175.9 | 171.9 | -179.2 | -177.3 | -178.1 | 170.3 | -158.8 |
| C11'-C10'-C9'-C8' | 172.9 | 173.3 | 172.8 | 175.2 | 174.8 | 174.2 | 174.6 | 174 | -179.7 |
| C10'-C9'-C8'-C7' | 178.4 | -23 | 178.3 | -160.3 | -175.1 | -180 | -179.4 | -170.2 | 174.9 |
| C9'-C8'-C7'-C6' | 176.3 | -171.6 | 176.3 | 177.6 | 178.5 | 177.2 | 176.8 | 177.3 | 178.6 |
| C8'-C7'-C6'-C5' (β2 ring) | -45.3 | 145.1 | -45.3 | -72.1 | -58.6 | -52.7 | -52.8 | -66 | -51.9 |
| RMSD | 0.436 | 0.706 | 0.436 |  |  |  |  |  |  |

\* Dihedral angles undergoing substantial change upon illumination are highlighted in grey.
\*RMSD was calculated with Pymol Align using PDB entry 4XB5 as a target selection

**Extended Data Table 2:**
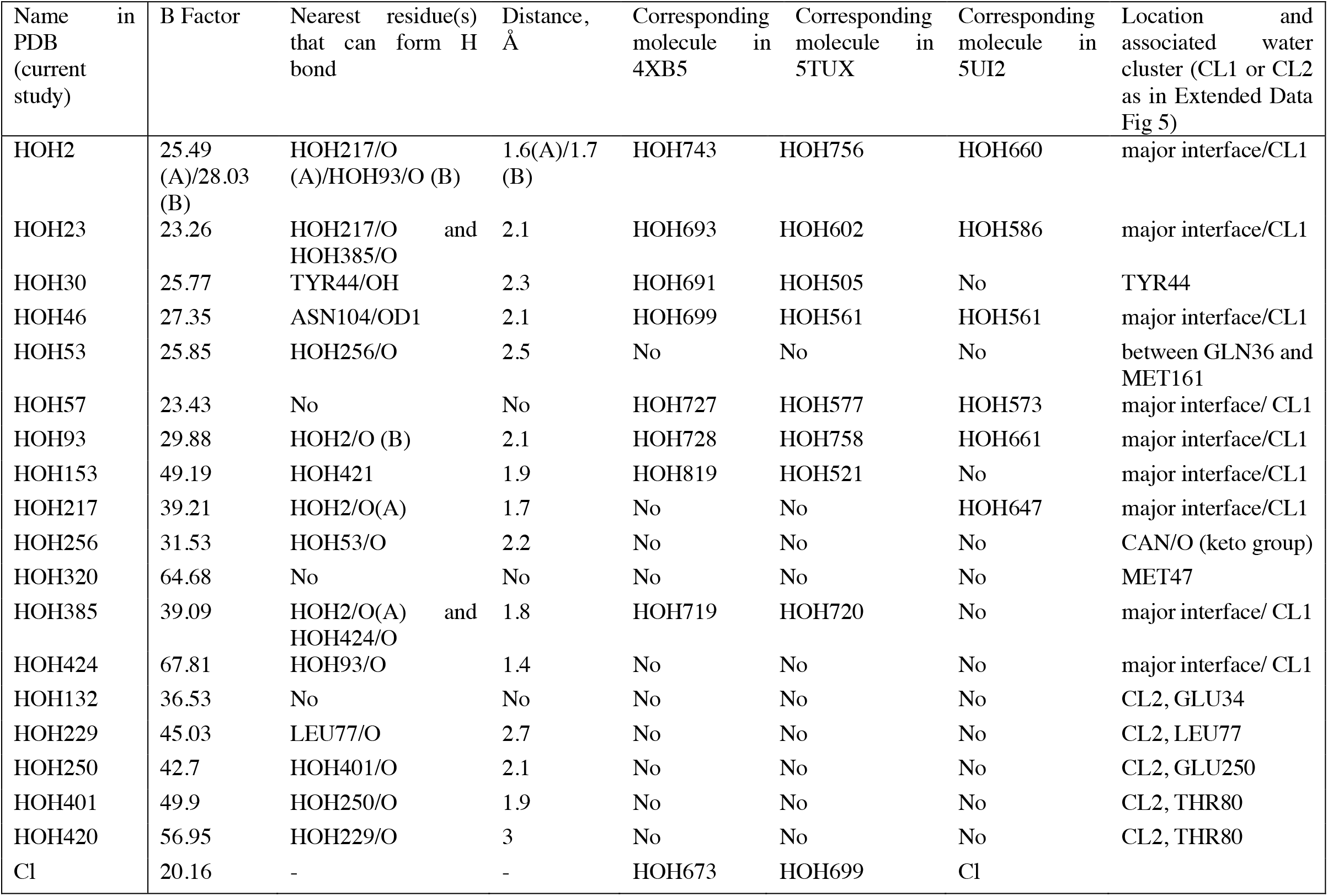
Cl^−^ and water molecules located within 6 Å of the carotenoid and in the newly identified water cluster (CL2). The molecules that are either conserved or found at the same location (within 0.5 Å) as in previously reported OCP structures (PDB entries 4XB5, 5TUX, 5UI2) are indicated.

| Name in PDB (current study) | B Factor | Nearest residue(s) that can form H bond | Distance, Å | Corresponding molecule in 4XB5 | Corresponding molecule in 5TUX | Corresponding molecule in 5UI2 | Location and associated water cluster (CL1 or CL2 as in Extended Data Fig 5) |
| --- | --- | --- | --- | --- | --- | --- | --- |
| HOH2 | 25.49 (A)/28.03 (B) | HOH217/O (A)/HOH93/O (B) | 1.6(A)/1.7 (B) | HOH743 | HOH756 | HOH660 | major interface/CL1 |
| HOH23 | 23.26 | HOH217/O and HOH385/O | 2.1 | HOH693 | HOH602 | HOH586 | major interface/CL1 |
| HOH30 | 25.77 | TYR44/OH | 2.3 | HOH691 | HOH505 | No | TYR44 |
| HOH46 | 27.35 | ASN104/OD1 | 2.1 | HOH699 | HOH561 | HOH561 | major interface/CL1 |
| HOH53 | 25.85 | HOH256/O | 2.5 | No | No | No | between GLN36 and MET161 |
| HOH57 | 23.43 | No | No | HOH727 | HOH577 | HOH573 | major interface/ CL1 |
| HOH93 | 29.88 | HOH2/O (B) | 2.1 | HOH728 | HOH758 | HOH661 | major interface/CL1 |
| HOH153 | 49.19 | HOH421 | 1.9 | HOH819 | HOH521 | No | major interface/CL1 |
| HOH217 | 39.21 | HOH2/O(A) | 1.7 | No | No | HOH647 | major interface/CL1 |
| HOH256 | 31.53 | HOH53/O | 2.2 | No | No | No | CAN/O (keto group) |
| HOH320 | 64.68 | No | No | No | No | No | MET47 |
| HOH385 | 39.09 | HOH2/O(A) and HOH424/O | 1.8 | HOH719 | HOH720 | No | major interface/ CL1 |
| HOH424 | 67.81 | HOH93/O | 1.4 | No | No | No | major interface/ CL1 |
| HOH132 | 36.53 | No | No | No | No | No | CL2, GLU34 |
| HOH229 | 45.03 | LEU77/O | 2.7 | No | No | No | CL2, LEU77 |
| HOH250 | 42.7 | HOH401/O | 2.1 | No | No | No | CL2, GLU250 |
| HOH401 | 49.9 | HOH250/O | 1.9 | No | No | No | CL2, THR80 |
| HOH420 | 56.95 | HOH229/O | 3 | No | No | No | CL2, THR80 |
| Cl | 20.16 | - | - | HOH673 | HOH699 | Cl |  |

**Extended Data Table 3:**
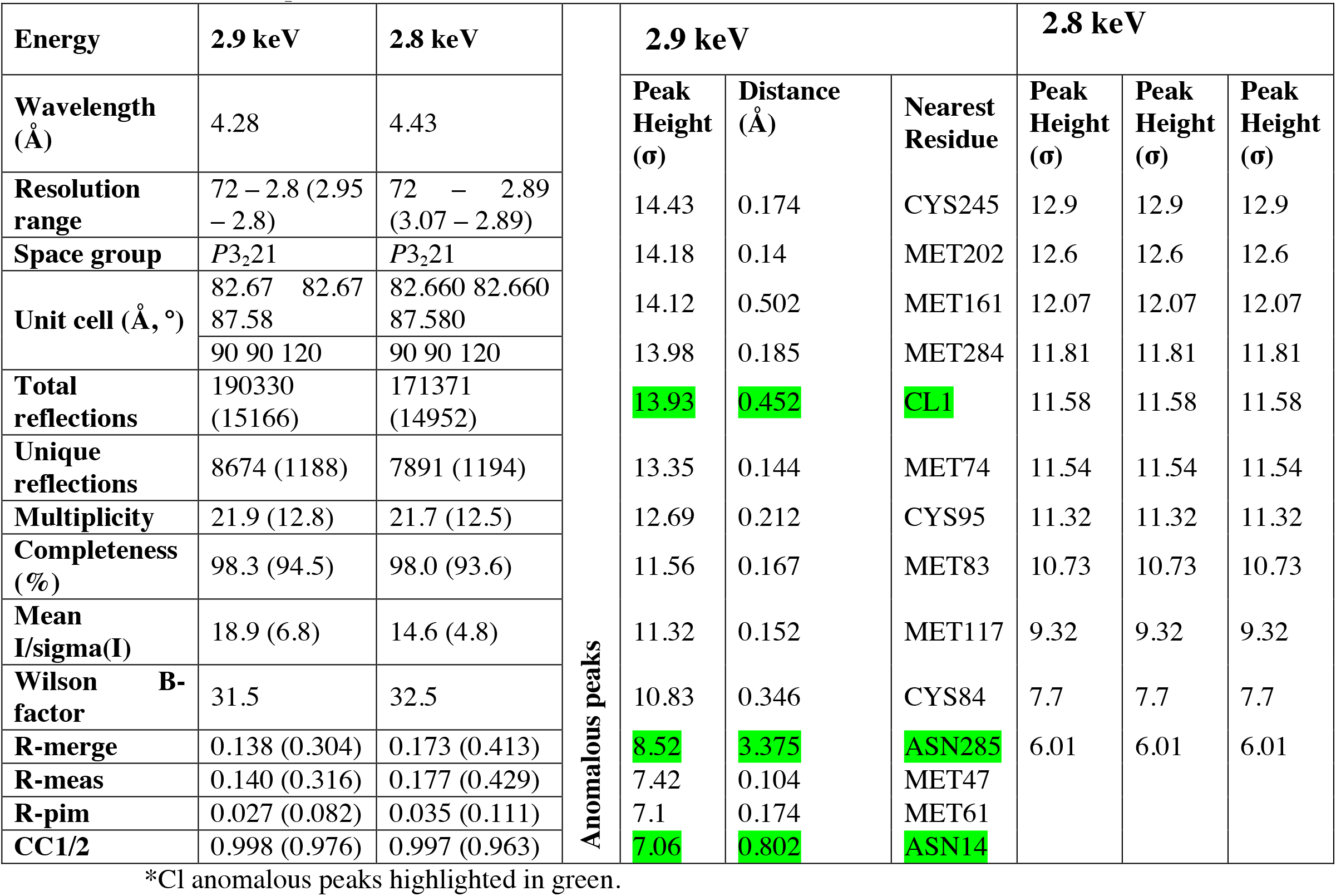
X-ray crystallography data collection, refinement statistics and anomalous peaks for OCP. Statistics for the highest-resolution shell are shown in parentheses.

**Extended Data Table 4:** Light-driven newly formed H-bonds in the NTD main chain. The H-bond list was obtained using the list_mc_hbonds.py script from Robert L. Campbell (http://pldserver1.biochem.queensu.ca/~rlc/work/pymol/). Water molecules were excluded from the calculations.

|  |  | Distances, Å |  |  |  |
| --- | --- | --- | --- | --- | --- |
| H-bond (inter subunit only) |  | I1 | I2 | I3 | I4 |
| A/ILE`25/N | A/PRO`22/O | 3.45 | 3.39 |  |  |
| A/ARG`27/N | A/THR`24/O | 3.53 | 3.38 | 3.26 |  |
| A/LEU`37/N | A/GLU`34/O | 3.35 | 3.32 |  | 3.29 |
| A/ALA`38/N | A/ASP`35/O | 3.35 | 3.34 |  |  |
| A/THR`52/N | A/THR`52/O | 3.09 |  |  |  |
| A/SER`60/N | A/GLY`57/O | 3.05 | 3.2 | 3.14 |  |
| A/GLU`65/N | A/LEU`63/O | 3.66 | 3.61 |  |  |
| A/LYS`69/N | A/ASN`66/O | 3.72 | 3.59 | 3.33 | 3.36 |
| A/CYS`84/N | A/GLN`81/O | 3.31 | 3.27 |  | 3.22 |
| A/ALA`87/N | A/CYS`84/O | 3.00 | 3.09 | 3.28 |  |
| A/TYR`98/N | A/CYS`95/O | 3.39 |  | 3.51 | 3.55 |
| A/LYS`106/N | A/PRO`103/O | 3.18 | 3.32 |  |  |
| A/TYR`111/N | A/GLY`108/O | 3.24 | 3.28 | 3.28 | 3.51 |
| A/LEU`113/N | A/TRP`110/O | 3.64 | 3.45 | 3.26 | 3.43 |
| A/ALA`140/N | A/ALA`137/O | 3.23 | 3.33 | 3.32 | 3.37 |

\* The pattern of newly formed and disrupted hydrogen bonds between intermediates illustrates the transient nature of the hydrogen bonding and agrees with our FTIR results (Fig 4). Out of 15 hydrogen bonds, most remain in the I1 and I2 structural intermediates, 7 do so in I1-I3 while 4 remain in all intermediates.

