## Supplementary material for "Light activation of Orange Carotenoid Protein reveals bicycle-pedal single-bond isomerization": SM

*Department of Life Sciences, Imperial College London, London SW7 2AZ, United Kingdom
#Diamond Light Source, United Kingdom

**Supplementary note 1: X-ray crystallography data collection and refinement statistics of OCP in the dark and four illuminated states I1-I4.**

| state | OCP$^{O}$ | | | OCP$^{O}$ | | | OCP$^{O}$ | | | I1 | | | I1 | | |
|---|---|---|---|---|---|---|---|---|---|---|---|---|---|---|---|
| dataset | 1 | | | 2 | | | 3 | | | 1 | | | 2 | | |
| | Overall | InnerShell | OuterShell | Overall | InnerShell | OuterShell | Overall | InnerShell | OuterShell | Overall | InnerShell | OuterShell | Overall | InnerShell | OuterShell |
| Low resolution limit | 71.48 | 71.48 | 1.46 | 30 | 30 | 1.35 | 87.29 | 87.29 | 1.4 | 71.49 | 71.49 | 1.35 | 37.39 | 37.39 | 1.5 |
| High resolution limit | 1.42 | 6.35 | 1.42 | 1.32 | 5.9 | 1.32 | 1.36 | 6.08 | 1.36 | 1.32 | 5.9 | 1.32 | 1.46 | 6.53 | 1.46 |
| Rmerge (within I+/I-) | 0.085 | 0.051 | 2.65 | 0.073 | 0.051 | 2.062 | 0.075 | 0.043 | 3.543 | 0.049 | 0.026 | 2.448 | 0.066 | 0.043 | 2.979 |
| Rmerge (all I+ and I-) | 0.086 | 0.052 | 2.735 | 0.074 | 0.053 | 2.15 | 0.076 | 0.043 | 3.624 | 0.05 | 0.027 | 2.528 | 0.067 | 0.044 | 3.104 |
| Rmeas (within I+ and I+/I-) | 0.089 | 0.053 | 2.805 | 0.076 | 0.054 | 2.239 | 0.079 | 0.045 | 3.73 | 0.051 | 0.028 | 2.661 | 0.069 | 0.046 | 3.162 |
| Rmeas (all I+ & I-) | 0.089 | 0.053 | 2.813 | 0.076 | 0.055 | 2.239 | 0.078 | 0.044 | 3.718 | 0.051 | 0.028 | 2.634 | 0.069 | 0.045 | 3.197 |
| Rpim (within I+/I-) | 0.027 | 0.016 | 0.915 | 0.024 | 0.017 | 0.864 | 0.025 | 0.014 | 1.165 | 0.016 | 0.009 | 1.02 | 0.022 | 0.014 | 1.054 |
| Rpim (all I+ & I-) | 0.02 | 0.012 | 0.656 | 0.017 | 0.013 | 0.618 | 0.018 | 0.01 | 0.83 | 0.012 | 0.007 | 0.721 | 0.016 | 0.011 | 0.762 |
| Rmerge in top intensity bin | 0.051 | - | - | 0.049 | - | - | 0.039 | - | - | 0.024 | - | - | 0.043 | - | - |
| Total number of observations | 1288728 | 14702 | 86669 | 1510774 | 15713 | 76625 | 1465081 | 17776 | 108032 | 1514258 | 18368 | 76396 | 1066826 | 13743 | 77009 |
| Total number unique | 65306 | 824 | 4777 | 81068 | 1001 | 5938 | 74252 | 939 | 5438 | 80954 | 1015 | 5917 | 60537 | 768 | 4419 |
| Mean((I)/sd(I)) | 15.3 | 50 | 1.1 | 17.1 | 48 | 1.2 | 17.1 | 63.4 | 1.1 | 23.8 | 95.8 | 1.2 | 19.1 | 57.3 | 1 |
| Mn(I) half-set correlation CC(1/2) | 0.999 | 0.999 | 0.526 | 0.999 | 0.999 | 0.535 | 1 | 0.999 | 0.53 | 1 | 0.999 | 0.509 | 0.999 | 0.999 | 0.448 |
| Completeness | 100 | 99.9 | 100 | 100 | 98.5 | 100 | 100 | 100 | 100 | 100 | 99.7 | 100 | 100 | 99.4 | 100 |
| Multiplicity | 19.7 | 17.8 | 18.1 | 18.6 | 15.7 | 12.9 | 19.7 | 18.9 | 19.9 | 18.7 | 18.1 | 12.9 | 17.6 | 17.9 | 17.4 |
| Mean(Chi^2) | 0.93 | 0.82 | 0.89 | 0.94 | 0.87 | 0.91 | 0.94 | 1.03 | 0.88 | 0.94 | 1.04 | 0.86 | 0.98 | 0.93 | 0.87 |
| Anomalous completeness | 100 | 99.6 | 100 | 100 | 96.8 | 100 | 100 | 100 | 100 | 100 | 99.5 | 99.9 | 99.8 | 99.9 | 100 |
| Anomalous multiplicity | 10.2 | 10.4 | 9.2 | 9.5 | 9 | 6.4 | 10.2 | 11 | 10.1 | 9.6 | 10.5 | 6.4 | 9 | 10.6 | 8.8 |
| DelAnom correlation between half-sets | -0.013 | 0.12 | 0.03 | 0.129 | 0.489 | -0.007 | -0.252 | -0.36 | -0.003 | -0.103 | 0.266 | -0.025 | -0.074 | -0.058 | -0.048 |
| Mid-Slope of Anom Normal Probability | 0.932 | - | - | 0.941 | - | - | 0.826 | - | - | 0.876 | - | - | 0.98 | - | - |

| state | I2 | | | I2 | | | I3 | | | I3 | | | I4 | | | I4 | | |
|---|---|---|---|---|---|---|---|---|---|---|---|---|---|---|---|---|---|---|
| dataset | 1 | | | 2 | | | 1 | | | 2 | | | 1 | | | 2 | | |
| | Overall | InnerShell | OuterShell | Overall | InnerShell | OuterShell | Overall | InnerShell | OuterShell | Overall | InnerShell | OuterShell | Overall | InnerShell | OuterShell | Overall | InnerShell | OuterShell |
| Low resolution limit | 29.07 | 29.07 | 1.46 | 55.29 | 55.29 | 1.37 | 55.5 | 55.5 | 1.44 | 87.07 | 87.07 | 1.32 | 87.01 | 87.01 | 1.5 | 29.86 | 29.86 | 1.41 |
| High resolution limit | 1.42 | 6.35 | 1.42 | 1.34 | 5.99 | 1.34 | 1.4 | 6.26 | 1.4 | 1.29 | 5.77 | 1.29 | 1.46 | 6.53 | 1.46 | 1.37 | 6.13 | 1.37 |
| Rmerge (within I+/I-) | 0.065 | 0.028 | 2.708 | 0.053 | 0.027 | 2.763 | 0.133 | 0.054 | 10.905 | 0.043 | 0.028 | 1.63 | 0.092 | 0.047 | 4.332 | 0.046 | 0.026 | 2.561 |
| Rmerge (all I+ and I-) | 0.067 | 0.03 | 2.79 | 0.054 | 0.028 | 2.847 | 0.135 | 0.055 | 11.077 | 0.044 | 0.029 | 1.725 | 0.093 | 0.047 | 4.419 | 0.046 | 0.027 | 2.654 |
| Rmeas (within I+ and I+/I-) | 0.069 | 0.03 | 2.867 | 0.056 | 0.029 | 2.971 | 0.14 | 0.057 | 11.51 | 0.045 | 0.03 | 1.82 | 0.096 | 0.049 | 4.564 | 0.048 | 0.027 | 2.74 |
| Rmeas (all I+ & I-) | 0.068 | 0.031 | 2.871 | 0.055 | 0.029 | 2.951 | 0.138 | 0.057 | 11.372 | 0.045 | 0.03 | 1.824 | 0.095 | 0.048 | 4.536 | 0.048 | 0.028 | 2.745 |
| Rpim (within I+/I-) | 0.021 | 0.01 | 0.94 | 0.017 | 0.009 | 1.079 | 0.044 | 0.018 | 3.645 | 0.014 | 0.009 | 0.813 | 0.03 | 0.015 | 1.43 | 0.015 | 0.009 | 0.966 |
| Rpim (all I+ & I-) | 0.015 | 0.008 | 0.673 | 0.013 | 0.007 | 0.767 | 0.031 | 0.013 | 2.554 | 0.01 | 0.007 | 0.583 | 0.021 | 0.011 | 1.017 | 0.011 | 0.007 | 0.692 |
| Rmerge in top intensity bin | 0.032 | - | - | 0.024 | - | - | 0.05 | - | - | 0.025 | - | - | 0.045 | - | - | 0.024 | - | |
| Total number of observations | 1271095 | 12807 | 86302 | 1474726 | 17861 | 81503 | 1364917 | 16392 | 100741 | 1564913 | 19356 | 60623 | 1182100 | 14432 | 87696 | 1378407 | 14775 | 80781 |
| Total number unique | 65092 | 806 | 4775 | 77412 | 973 | 5680 | 68765 | 865 | 5049 | 86400 | 1081 | 6302 | 59958 | 773 | 4396 | 71802 | 900 | 5231 |
| Mean((I)/sd(I)) | 19.1 | 67.3 | 1.1 | 23 | 100.1 | 1 | 14.4 | 45.5 | 1.4 | 25.6 | 95.6 | 1.2 | 16.5 | 53.3 | 1.4 | 24.6 | 94 | 1 |
| Mn(I) half-set correlation CC(1/2) | 1 | 0.999 | 0.52 | 1 | 1 | 0.468 | 0.998 | 0.985 | 0.739 | 1 | 0.999 | 0.561 | 0.999 | 0.999 | 0.688 | 1 | 1 | 0.533 |
| Completeness | 100 | 98.5 | 100 | 100 | 99.8 | 100 | 100 | 99.6 | 100 | 100 | 99.4 | 100 | 100 | 100 | 100 | 100 | 98.6 | 100 |
| Multiplicity | 19.5 | 15.9 | 18.1 | 19.1 | 18.4 | 14.3 | 19.8 | 19 | 20 | 18.1 | 17.9 | 9.6 | 19.7 | 18.7 | 19.9 | 19.2 | 16.4 | 15.4 |
| Mean(Chi^2) | 0.94 | 0.64 | 0.95 | 0.91 | 1.1 | 0.84 | 0.97 | 0.92 | 1.01 | 0.89 | 1.07 | 0.74 | 0.95 | 1.01 | 0.83 | 0.88 | 0.88 | 0.76 |
| Anomalous completeness | 100 | 98.5 | 100 | 100 | 99.9 | 100 | 99.9 | 99.7 | 99.7 | 100 | 99.3 | 99.6 | 100 | 100 | 99.9 | 100 | 98.6 | 100 |
| Anomalous multiplicity | 10 | 9 | 9.1 | 9.8 | 10.8 | 7.3 | 10.2 | 11.1 | 10.1 | 9.3 | 10.3 | 4.8 | 10.2 | 11 | 10.2 | 9.9 | 9.5 | 7.8 |
| DelAnom correlation between half-sets | 0.131 | 0.728 | -0.025 | -0.034 | 0.067 | -0.029 | -0.154 | -0.243 | 0.004 | -0.118 | -0.127 | 0.019 | -0.323 | -0.47 | -0.005 | -0.104 | 0.275 | -0.006 |
| Mid-Slope of Anom Normal Probability | 0.912 | - | - | 0.899 | - | - | 0.797 | - | - | 0.893 | - | - | 0.782 | - | - | 0.882 | - | - |

| | OCP$^{O}$ 1 | OCP$^{O}$ 2 | OCP$^{O}$ 3 | I1 1 | I1 2 | I2 1 | I2 2 | I3 1 | I3 2 | I4 1 | I4 2 |
|---|---|---|---|---|---|---|---|---|---|---|---|
| **Average unit cell** | 82.54 82.54 87.35 90 90 120 | 82.54 82.54 87.36 90 90 120 | 82.59 82.59 87.29 90 90 120 | 82.55 82.55 87.21 90 90 120 | 82.68 82.68 87.66 90 90 120 | 82.48 82.48 87.21 90 90 120 | 82.52 82.52 87.26 90 90 120 | 82.81 82.81 87.66 90 90 120 | 82.50 82.50 87.07 90 90 120 | 82.58 82.58 87.01 90.00 90.00 120.00 | 82.44 82.44 86.63 90.00 90.00 120.00 |
| **Space group** | *P*3₂21 | *P*3₂21 | *P*3₂21 | *P*3₂21 | *P*3₂21 | *P*3₂21 | *P*3₂21 | *P*3₂21 | *P*3₂21 | *P*3₂21 | *P*3₂21 |

| **Refinement** | OCP$^{O}$ 1 | OCP$^{O}$ 2 | OCP$^{O}$ 3 | I1 1 | I1 2 | I2 1 | I2 2 | I3 1 | I3 2 | I4 1 | I4 2 |
|---|---|---|---|---|---|---|---|---|---|---|---|
| Resolution | 55.39-1.43 | 30.02-1.32 | 55.40-1.36 | 55.35-1.32 | 37.42-1.46 | 29.09-1.42 | 55.35-1.34 | 55.36-1.39 | 71.55-1.29 | 71.62-1.46 | 29.89-1.37 |
| No. reflections all/free | 63888 / 3134 | 81019 / 3975 | 74207 / 3699 | 80887 / 3975 | 60499 / 2818 | 65042 / 3188 | 77359 / 3895 | 69455 / 3445 | 86348 / 4173 | 59930 / 2794 | 71743 / 3474 |
| R-factor/R-free | 0.168 / 0.203 | 0.170 / 0.192 | 0.172 / 0.196 | 0.167 / 0.192 | 0.164 / 0.192 | 0.168 / 0.200 | 0.170 / 0.192 | 0.1656 / 0.189 | 0.171 / 0.1954 | 0.163 / 0.190 | 0.175 / 0.217 |
| RMS Deviations | | | | | | | | | | | |
| Bonds | 0.0121 | 0.0148 | 0.0142 | 0.015 | 0.0132 | 0.0131 | 0.0147 | 0.01485 | 0.0149 | 0.0133 | 0.0133 |
| Angles | 1.871 | 2.032 | 1.988 | 2.054 | 1.893 | 1.91 | 2.012 | 2.015 | 2.028 | 1.98 | 1.946 |
| Chain mean B | | | | | | | | | | | |
| (No. atoms) | | | | | | | | | | | |
| AAA | 25.5( 5563 ) | 23.5( 5671 ) | 24.4( 5563 ) | 24.0( 5668 ) | 29.7( 5626 ) | 26.4( 5563 ) | 24.8( 5563 ) | 27.6( 5563 ) | 23.7( 5587 ) | 26.7( 5680 ) | 27.2( 5671 ) |
| AaA | 18.8( 94 ) | 16.6( 94 ) | 17.5( 94 ) | 17.8( 94 ) | 22.9( 94 ) | 19.5( 94 ) | 17.9( 94 ) | 20.9( 94 ) | 17.3( 94 ) | 19.7( 94 ) | 19.6( 94 ) |
| AbA | 27.0( 12 ) | 25.1( 12 ) | 26.9( 12 ) | 26.6( 12 ) | 32.8( 12 ) | 28.5( 12 ) | 26.7( 12 ) | 30.0( 12 ) | 25.3( 12 ) | 29.8( 12 ) | 29.0( 12 ) |
| AcA | 70.5( 7 ) | 72.0( 7 ) | 65.7( 7 ) | 78.2( 7 ) | 82.0( 7 ) | 72.1( 7 ) | 68.3( 7 ) | 69.3( 7 ) | 64.2( 7 ) | 72.5( 7 ) | 68.6( 7 ) |
| BBB | 41.5( 449 ) | 39.7( 437 ) | 41.9( 449 ) | 41.7( 494 ) | 46.5( 449 ) | 42.5( 449 ) | 41.3( 49 ) | 44.4( 449 ) | 40.4( 471 ) | 45.7( 543 ) | 45.4( 512 ) |
| CCC | 20.9( 1 ) | 18.7( 1 ) | 20.1( 1 ) | 19.7( 1 ) | 25.1( 1 ) | 21.3( 1 ) | 19.9( 1 ) | 22.9( 1 ) | 19.4( 1 ) | 22.1( 1 ) | 22.9( 1 ) |
| **Ramachandran** | | | | | | | | | | | |
| In preferred regions | 192 (94.58%) | 191 (95.5%) | 193 (95.07%) | 196 (96.17%) | 192 (95.52%) | 192 (94.58%) | 193 (95.57%) | 194 (95.57%) | 194 (95.57%) | 186 (93.47%) | 188 (94.0%) |
| in allowed regions | 11 (5.42%) | 9 (4.5%) | 10 (4.93%) | 7(3.83%) | 9(4.48%) | 11 (5.42%) | 9 (4.43%) | 9 (4.43%) | 9 (4.43%) | 13(6.53%) | 12(6.0%) |
| Outliers | 0 (0%) | 0 (0%) | 0 (0%) | 0 (0%) | 0 (0%) | 0 (0%) | 0 (0%) | 0 (0%) | 0 (0%) | 0 (0%) | 0 (0%) |

### Supplementary note 2: Monitoring the OCP photocycle and primary photoproduct in solution by UV-VIS spectroscopy

To confirm that the C9'-C8' *cis* and C7'-C6' *trans* (bicycle-pedal isomer) of CAN in I1 thermally relaxes back to C9'-C8' *trans* and C7'-C6' *cis* after 2 min of illumination and rule out a light-driven process, we collected time-dependent UV-VIS spectra upon 40 s of illumination followed by 5 min of darkness (Extended Data Fig 4 a). Similar to the experiment from Fig. 4a, all the spectra collected within 5 min 40 s were fit simultaneously using global fit approach[1–3] . In these experimental conditions relaxation of the bicycle-pedal CAN isomer, if light driven, should not exhibit a spectral characteristic of EAS3 observed in Fig 4a upon continuous illumination.

The results of the global fitting of the dataset highlight four EAS components (Extended Data Fig 4 a). The EAS2 component from Extended Data Fig 4 a (red solid line), like its counterpart in Fig 4a, due the spectral similarity of the two spectra, can with confidence be assigned to the formation of the bicycle-pedal isomer like in Fig 4 a. The EAS3 component (Extended Data Fig 4 a ,dotted line) shows a 25 nm red-shifted absorption maximum similar to that observed in Fig. 4a, thereby confirming that the bicycle-pedal isomer thermally relaxes to its "dark " C9'-C8' *trans* and C7'-C6' *cis* conformation in 2 min to yield a new $OCP^R$-like intermediate. The steady-state EAS4 component represents recovery of the $OCP^R$-like intermediate state to $OCP^O$, similar to the $OCP^R$-$OCP^O$ conversion occurring in darkness.

To demonstrate the light intensity dependence of both the rate of accumulation of the primary product and photocycle intermediates, we monitored absorption changes of OCP at different power densities and two characteristic wavelengths: 470 nm ($OCP^O$/Isom) and 550 nm ($OCP^R$-like, $OCP^R$) (Extended Data Fig 4 b-c). The results show that the rates of absorption changes collected at both wavelengths are proportional to illumination intensities

### Supplementary note 3: Light-driven salt bridge formation and $Cl^-$ displacement in OCP

As can be seen in Fig 3b, the position of R155 is modulated by HOH41 and HOH46 via their H-bonding with R155. In I1, the strength of HOH41-R155 and HOH46-R155 H-bonds increases as their hydrogen donor – hydrogen acceptor distances decrease by 0.3 Å and 0.8 Å respectively. This H-bond rearrangement is controlled by the chloride ion which is located close to the domain interface in the CTD where it coordinates four water molecules (HOH13, HOH46, HOH41 and HOH23) within a 3.0-4.0 Å radius through ion-dipole interactions (Extended Data Fig 6 a).

In the I1-I3 intermediates, a small displacement of the chloride ion by 0.2 Å coincides with small displacements of all four water molecules it is hydrogen-bonded to through ion-dipole forces (Extended Data Fig 6 a, Fig. 3b).

After 2-5 min of illumination (I2, I3, Fig 3c), the HOH46-R155 distance remains the same as in I1 whereas the HOH41-R155 distance reverts back to its original value in $OCP^O$. After 10 min the HOH46-R155 hydrogen bond strength turns from moderate to strong since its distance decreases down to 2.5 Å. Unlike water molecules, all CTD amino acids in I1 located within a 4 Å radius from $Cl^-$ (W277, P276, T275 and L248) remain in their position upon illumination, with the exception of a marginal (<<0.2 Å) displacement of P276.

In I2-I3 these amino-acids are displaced by 0.2 Å, coinciding with the movement of other CTD amino acids located at the domain interface. In I4 all amino acids within 4 Å from $Cl^-$ undergo a translocation by 0.5 Å with $Cl^-$ moving in the same direction. Overall, the movement of the chloride ion observed in the first 5 min of illumination is mainly caused by the H-bond rearrangement in the water channel (Cl1, Extended Data Fig 5, Extended Data Fig 6 a), while in I4 it is mainly caused by translocation of the CTD away from NTD (Extended Data Fig 6 b).

### Supplementary note 4: water molecules in FTIR data

It should be kept in mind that the positive and negative features of the EAS FTIR spectra (Fig 4c,d and Extended Data Fig 4 f) are the result of broad overlapping contributions from positive and negative spectral signatures as well as one or more vibration modes giving a signal at the same wavenumber. Since it is not possible to disentangle such overlapping contributions, it is difficult to assign spectral features solely to a single vibration process.
The most complicated spectral features are in the 1600−1645 $cm^{-1}$ region, where the spectral differences between the $H_2O$ and the $D_2O$ spectra are not easily interpretable by simple H/D substitution. For instance, the ~1600−1645 $cm^{-1}$ band (in $H_2O$) observed between 40-120 s after the illumination onset (EAS2-3, Extended Data Fig 4 f), completely disappears in the presence of $D_2O$ (Fig 4c). Similar FTIR signature has been observed previously in OCP where it was tentatively assigned to $H_2O$ vibration modes[4]. Indeed, only the HOH symmetric mode vibration (1643 $cm^{-1}$) contributes in the observed region (1700 $cm^{-1}$-1500 $cm^{-1}$) whereas the DOD symmetric vibration band lies at 1210 $cm^{-1}$, (Extended Data Fig 4 g). This therefore suggests that the ~1600−1645 $cm^{-1}$ shoulder could be assigned to vibrational motions of HOH. In support of this assignment, we observed that the presence of the 1600-1640 $cm^{-1}$ band in the $H_2O$ data correlates with the light-driven displacement of water molecules observed in the crystallographic data on the same time scale (Extended Data Fig 5- I2 and I3). The rearrangement of water molecules affects the two water clusters CL1 and CL2 each centred on HOH46 and HOH152 respectively. The strongest DED signals are observed in CL1, which is located on the interface between the two domains (Extended Data Fig 5, Extended Data Fig 4 f).
